# Transcriptomically unique endolysosomal and homeostatic microglia populations in Alzheimer’s disease and aged human brain

**DOI:** 10.1101/2021.10.25.465802

**Authors:** Katherine E. Prater, Kevin J. Green, Sainath Mamde, Wei Sun, Alexandra Cochoit, Carole L. Smith, Kenneth L. Chiou, Laura Heath, Shannon E. Rose, Jesse Wiley, C. Dirk Keene, Ronald Y. Kwon, Noah Snyder-Mackler, Elizabeth E. Blue, Benjamin Logsdon, Jessica E. Young, Ali Shojaie, Gwenn A. Garden, Suman Jayadev

## Abstract

Microglia contribute to Alzheimer’s Disease (AD) progression and are candidate therapeutic targets. Human microglia exhibit an array of transcriptional phenotypes implying that accurate manipulation of microglial function will require clarity of their molecular states and context dependent regulation. To increase the number of microglia analyzed per subject we employed fluorescence activated nuclei sorting prior to single-nucleus RNA-seq on human prefrontal cortices. We observed microglia phenotypes previously unrecognized in human brain gene expression studies and mapped their transcriptomic relationships by trajectory inference. Three clusters were enriched for endolysosomal pathways, one of which showed differential expression of AD GWAS genes in addition to genes implicated in nucleic acid detection and interferon signaling. Analysis of the “homeostatic” microglia cluster revealed a uniquely AD subcluster. Our study demonstrates the value of deeply profiling microglia to explore the biological implications of microglia transcriptomic diversity.

## Introduction

As the most common form of aging-related cognitive decline, Alzheimer’s Disease (AD) affects millions of individuals worldwide. AD is pathologically characterized by the presence of extracellular amyloid-beta plaques, neuronal intracellular neurofibrillary tangles, and neuroinflammation. Microglia are the resident innate immune cells of the brain, and contribute to the neuroinflammatory processes hypothesized to promote AD pathophysiology^1–9^. In AD brain, microglia release inflammatory factors with non-cell-autonomous effects, lose protective homeostatic function, and initiate aberrant phagocytosis of synapses and neurons, all of which influence the pathophysiology behind cognitive decline^5,9–11^. As such, microglia inflammatory responses have potential as future therapeutic targets. Yet large gaps remain in our understanding of microglia responses in AD brain.

In line with their known diversity of function, microglia assume a spectrum of phenotypes. These can be differentiated by morphology, physiology and gene or protein expression patterns, though most of this understanding has come from model systems^6,12–16^. Less is known about the heterogeneity of microglia states within the adult human brain in health and neurodegenerative disease. Single cell and single nuclei RNA sequencing (snRNAseq) studies of fresh and frozen human cortical tissue have revealed the diversity of microglia phenotypes in the context of AD and other brain pathologies^17–24^. Distinguishing transcriptomically distinct clusters enables the identification of candidate genetic and epigenetic factors regulating specific phenotypes which can be leveraged for precision therapeutics approaches. However standard snRNAseq methods have been restricted by the small number of microglia assayed per individual limiting the potential to map the full range of microglial subpopulations. We posited that cellular processes and regulatory factors contributing to “responsive” microglia profiles would be uncovered by improved resolution of microglia phenotypes in human brain. To enhance characterization of the changes occurring in microglia phenotypes in AD, we employed a microglia enrichment technique for single nuclei RNAseq. We generated microglia transcriptional profiles from a cohort of 22 individuals, annotated microglia clusters with plausible biological roles and identified differences between AD and control individuals within microglial phenotypes. The depth of our dataset also allowed us to investigate the diversity of subclusters within the microglia cluster typically annotated as “homeostatic”. We found a homeostatic marker enriched population unique to AD which may hold clues to early or subtle microglial changes attributable to AD pathology. In addition to homeostatic and inflammatory phenotypes described previously, we uncovered microglial subpopulations with distinct transcriptomic profiles that provide new avenues for hypotheses testing in future studies of AD mechanisms.

## Results

### Fluorescence-activated nuclei sorting for PU.1 expression enriches microglia nuclei 20-fold

We enriched the nuclei isolated from post-mortem human brain for microglia using fluorescence-activated nuclei sorting (FANS) for expression of the myeloid specific transcription factor (PU.1; Figure S1). FANS plots of the PU.1 staining and controls are demonstrated in Figure S1A,B). To confirm that PU.1 FANS was effective, we isolated and sequenced nuclei with and without PU.1 FANS (N=4). We analyzed similar numbers of total nuclei in the unsorted (46,085; Figure S1C) and PU.1 sorted (41,488; Figure S1D) datasets. The PU.1 sorted dataset contained 20X more microglia nuclei defined by high expression of *C3, CD74, C1QB, CX3CR1*, and *SPI1* (23,310 microglia nuclei) than the unsorted dataset (1,032 microglia nuclei; Figure S1C,D). The microglia nuclei observed in the PU.1 sorted dataset also demonstrate further complexity as evidenced by more microglia clusters (Figure S1D).

We next applied PU.1 FANS to a cohort of 22 individuals (Figure 1A). After PU.1 FANS, samples retain a variety of non-myeloid cell types (Figure 1B) while providing clear resolution of clusters demonstrating distinct microglia gene expression patterns (63% of the nuclei; Figure 1C). This dataset is the largest microglia per sample dataset generated thus far, including compared to other published datasets that have used alternative enrichment techniques. Gene expression of cell type marker genes demonstrates that the clusters identified as microglia (1, 2, 3, 5, 16, and 17) in the dataset have high expression of microglia markers and do not express canonical marker genes of other cell types (Figure 1D). Detailed gene expression plots of both microglia markers (Figure S2), and astrocyte or peripheral monocyte markers (Figure S3) demonstrate high expression of microglia genes in the microglia subset dataset across all subclusters and the lack of other cell type and peripheral markers.

**Figure 1.**
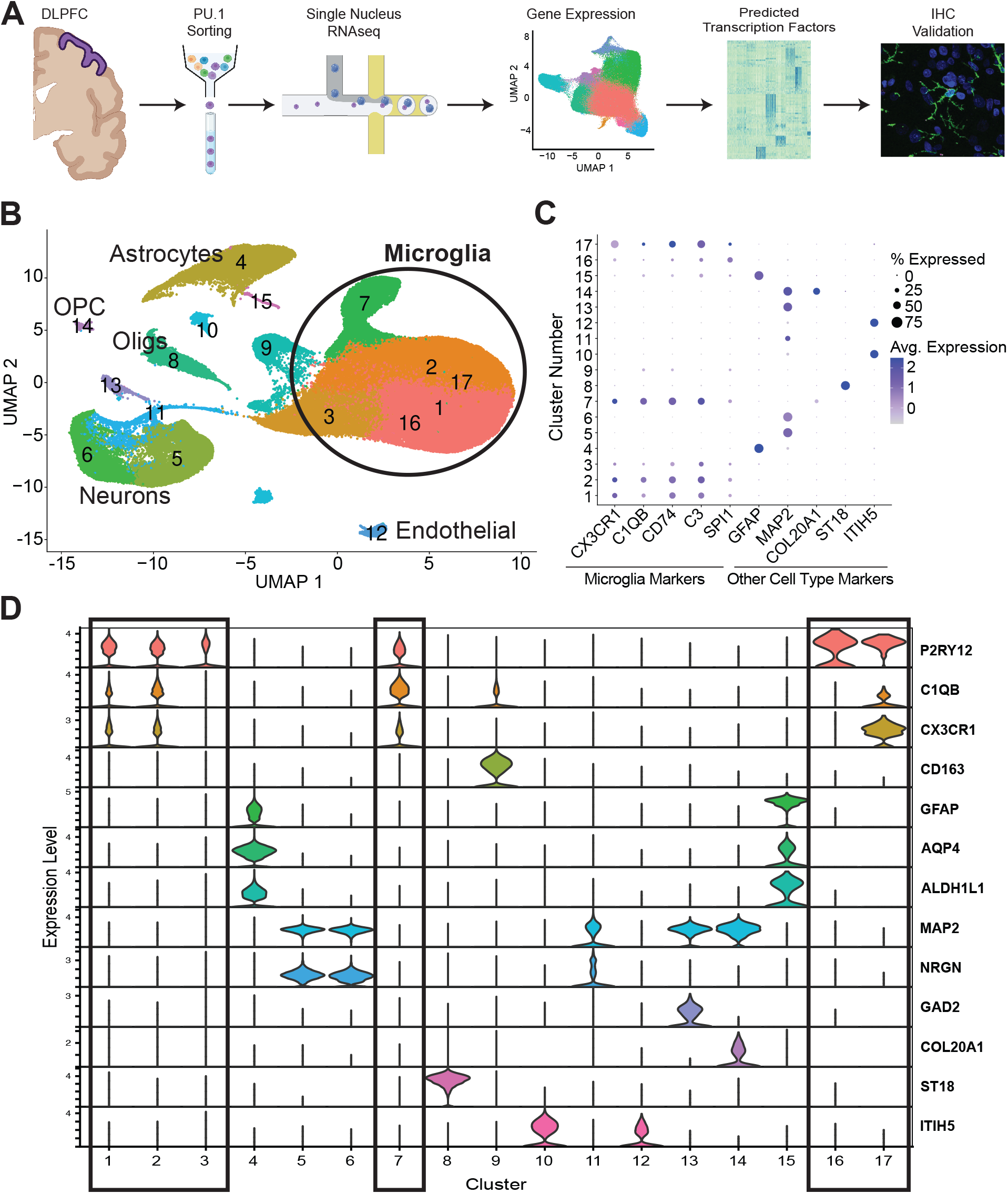
PU.1 enrichment yields a large dataset of microglia nuclei. **A)** Experimental design of 22 post-mortem human prefrontal cortices. (Created in part with BioRender.com) **B)** UMAP of the 22 subject dataset demonstrates that while other cell types including neurons, astrocytes, oligodendrocytes (Oligs) and their progenitors (OPC) as well as endothelial cells are present, six clusters including the three largest are composed of microglia nuclei. **C)** Representative cell type marker genes (x-axis) with the percent of nuclei that express a gene (size of dot) in each cluster (distributed along Y axis) and the average expression level (color intensity) are shown for microglia (*CX3CR1, C1QB, CD74*, and *C3*), astrocytes (*GFAP*), neurons (*MAP2*), OPCs (*COL20A1*), oligodendrocytes (*ST18*), and endothelial cells (*ITIH5*) for each cluster. **D)** Gene expression of a wider set of cell type marker genes demonstrate that clusters 1, 2, 3, 7, 16, and 17 are composed of microglia.

### Complexity of microglia states

The initial dataset consisted of 205,226 nuclei, with 200,948 nuclei (98%) passing QC and doublet removal. Of those, 127,371 were identified as microglia (Figure 1D, Figure S2), with an average of 5,790 nuclei per individual. Cluster analysis of nuclei with microglia-like expression identified 10 clusters (Figure 2A) characterized by differentially expressed genes (DEGs) comparing the cluster to all other nuclei (Figure 2B). Using gene set enrichment analysis (GSEA), we determined the enrichment of biological pathways in each cluster (Figure 2C). There was little to no overlap in the DEGs defining each cluster or the biological pathways identified by GSEA supporting the uniqueness of each cluster.

**Figure 2.**
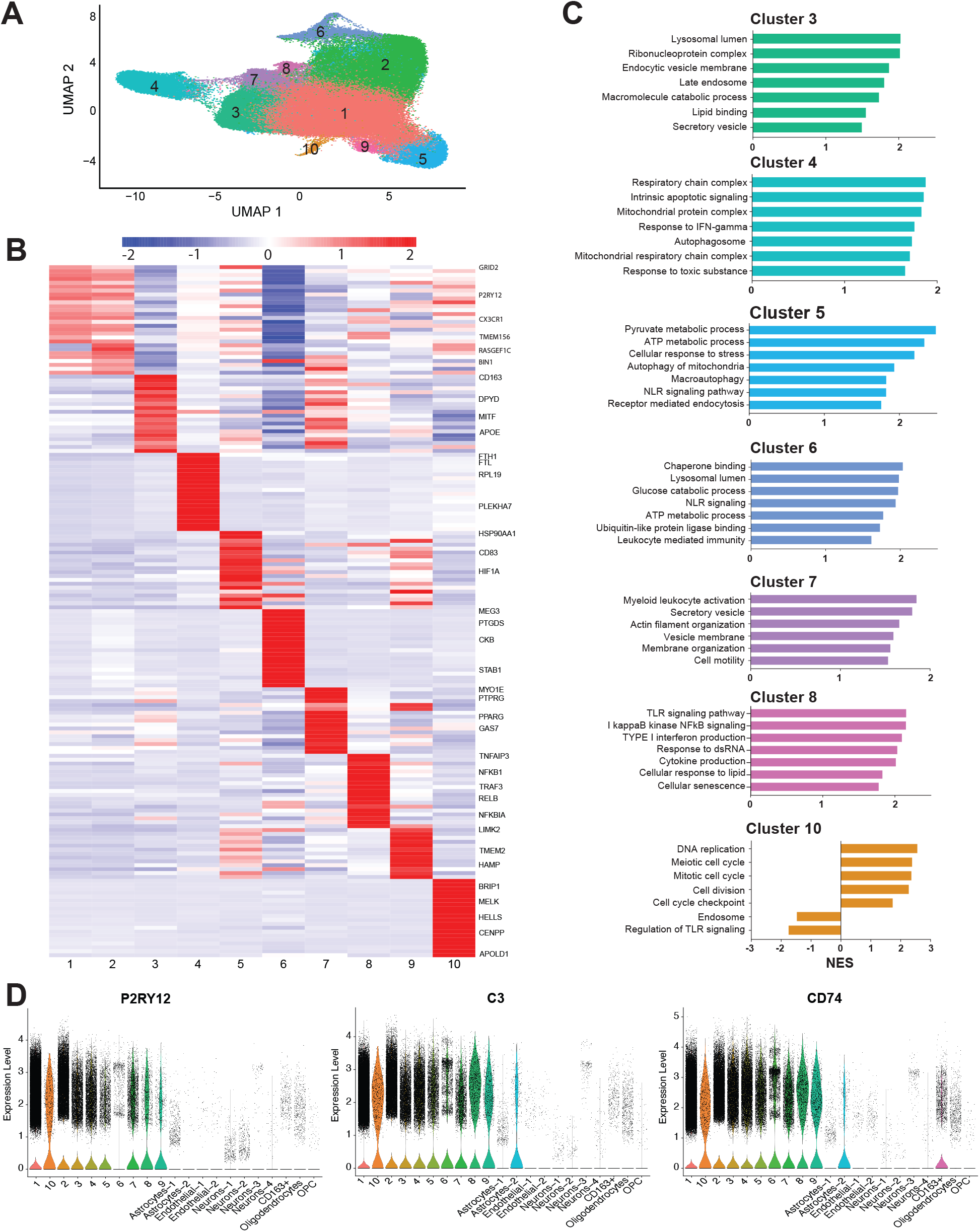
Microglia states have diverse gene expression and biological pathway correlates. **A)** UMAP. **B)** Top 25 genes from each cluster. **C)** GSEA analysis suggests distinct biological pathways. **D)** Canonical microglia marker gene expression in the microglia dataset versus cell types sorted during PU.1 enrichment. (NES = Normalized Enrichment Score)

First, we found clusters with annotations similar to microglia phenotypes previously described in human brain. We identified Cluster 1, the largest cluster, as the cluster enriched with homeostatic markers including high expression of *CX3CR1* and *P2RY12*^15,19–21^. We abbreviate this homeostatic marker expressing cluster as “HM”. Although different in their gene expression, HM and Cluster 2 may have similar biological function (Figure 2B). HM was thus established as the basis for comparison to assess DEGs for other clusters, replicating the approach employed in previous publications^17,20,21^ (Table S2). Cluster 4 is enriched for pathways involved in apoptosis, response to interferon-gamma (IFNγ), and mitochondrial and respiratory functions (Figure 2C) including Alzheimer, Parkinson, and Huntington Disease KEGG pathways. The most highly differentially expressed gene in this cluster is *FTL*. Taken together, the profile of Cluster 4 is suggestive of a degenerative or dystrophic phenotype^21^. Cluster 7 is characterized by expression of genes involved in migration and motility (Figure 2B). Pathways enriched in Cluster 7 also indicate these cells are motile, with changes to cytoskeleton and membrane that indicate movement of processes or the cell itself (Figure 2C). Cluster 8 featured a canonical inflammatory phenotype with expression of classic inflammatory activation genes including *NFκB1, RelB*, and *IL1β*,; Figure 3B)^1,25,26^. GSEA revealed this cluster was enriched in NFκB signaling, Interferon signaling, Toll-like receptor and RIG-1 mediated signaling pathways indicating downstream effector inflammatory responses to stimuli (Figure 2C). Cluster 9 is defined by genes and pathways involved in iron homeostasis and cytokine production (Figure 2B). This profile is consistent with senescent microglia^27^. Cluster 10 is defined by expression of genes involved in cell cycle regulation and DNA repair^28,29^ (Figure 2B). The pathways enriched in Cluster 10 confirm the relative increase of genes involved in cell cycle processes and a decrease of endosome and cytokine processing genes in these microglia (Figure 2C). As expected for microglia of varying activated and non-activated phenotypes, genes such as P2RY12 varied in expression across the 10 microglia subclusters, while microglia genes such as C3 and CD74 had more similar representation across the microglia subclusters (Figure 2D).

**Figure 3.**
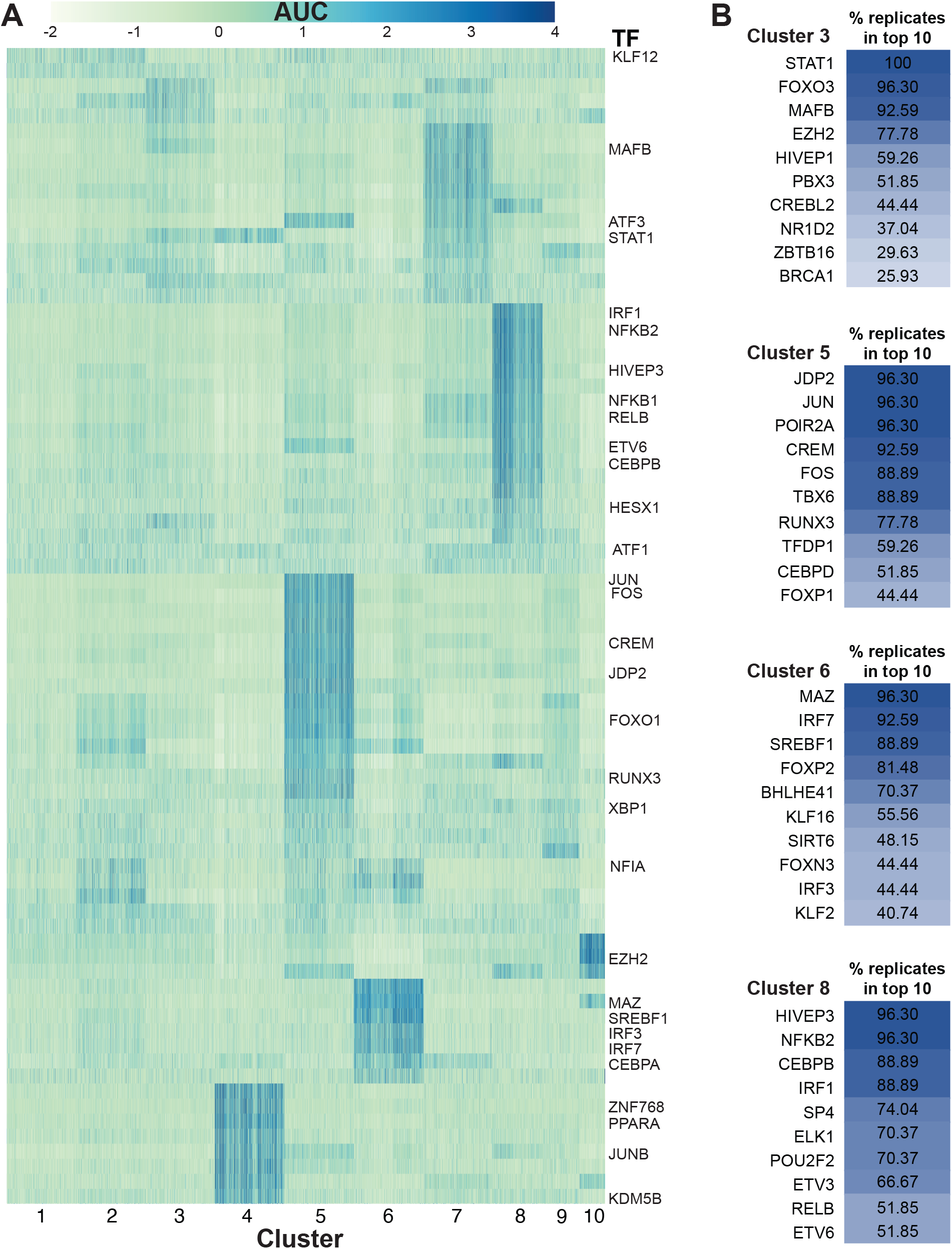
Transcription factor regulatory networks are specific to microglia phenotypes. **A)** Heatmap of area under the curve (AUC) of the transcription factor (TF) regulon activity in clusters. **B)** Percentage of instances where transcription factor regulons occured in the top ten regulon specificity scores per cluster (out of 27 permutations).

Next, we found three clusters, Clusters 3, 5 and 6, which have not been previously described in human brain. These clusters were distinguished by their significantly increased endolysosomal network (ELN) gene expression and enrichment for ELN pathway signatures relative to HM microglia. We therefore annotate them collectively as ELN in this report. Cluster 3 was defined by genes implicated in aggregate protein internalization (Figure 2B)^15,16,19–21,30^ phagocytosis, and vesicle mediated transport ^26^. Pathways enriched in Cluster 3 include endosome and lysosome pathways as well as catabolism and lipid binding, but no inflammatory processes (Figure 2C). Genes involved in glycolysis have lower expression in Cluster 3 differentiating it from the two other ELN clusters, suggesting these cells have not undergone the metabolic switch to glycolysis observed in the microglia inflammatory phenotype^31^. Cluster 5 and 6 displayed an ELN signature, though also appeared metabolically active with distinct inflammatory characteristics. Cluster 5 had increased expression of *HSP90AA1, HIF1A*, and *BAG3* in addition to other heat shock protein (HSP) genes (Figure 2B), suggesting these cells are responding to external stress. Genes driving glycolysis were also higher in Cluster 5 when compared to HM, possibly reflecting a switch to glycolysis in these cells^31^. The pathways enriched in this cluster indicate it is active in endocytosis, autophagy, and mitophagy (Figure 3C). Cluster 6 is characterized by metabolic activity genes and stress response pathways similar to Cluster 5 (Fig 2C), though with an additional component of interferon signaling suggested by significantly higher levels of *IRF3, IRF5*, and *IRF7*. Cluster 6 also showed increased expression of cytosolic DNA/RNA recognition and antiviral genes including *IFIT2, IFIT3,TRIM22* (34440633) as well as the pattern recognition receptor *CARD9*, and mediators of the NLRP3 inflammasome^25,26,32–35^. Of note, unique to Cluster 6, we found that the DNA repair genes^28^ *ATM*, and *RNASEH2B* were lower in expression. While *IL1β* was increased in Cluster 6 compared to HM, even higher levels of *IL1β* and expression of other inflammatory effector molecules such as *NFkB* were found in Cluster 8, the more canonical “inflammatory” cluster. Pathways enriched in Cluster 6 also support upstream inflammatory responses to danger associated molecular pattern stimuli such as enrichment in NOD-like receptor signaling. We also used alternative methodology to identify biological pathways and the links between them, providing validation for the endolysosomal network functions of clusters 3, 5, and 6 (Figure S4). Supplemental tables containing the genes driving the presence of each node in the network are available on Synapse.

Despite the role of *APOE* in risk and progression of AD^36^, to date no studies have defined microglia states in individuals with a specific *APOE* genotype. Since the majority (13/22) of our sample were homozygous for the *APOE* ε3 allele, we generated a subset of our dataset that consisted entirely of *APOE* ε3/ε3 individuals (7 controls and 6 AD pathology; 75,018 microglia nuclei). After re-normalizing and re-clustering, we identified 9 clusters of microglia (Figure S5A). These clusters were defined by genes similar to those that defined the clusters in the Mixed *APOE* genotype dataset (Figure S5B). We found that in most clusters the DEGs were very similar (∼60% or higher match) to those in the Mixed *APOE* dataset (Figure S5C). The HM, neurodegenerative, inflammatory, cell cycle, and endolysosomal clusters were similar to those of the Mixed *APOE* cohort (Figure S5C). This suggests that the presence of multiple distinct ELN microglia clusters is common in the human brain even when controlling for *APOE* genotype.

### Transcription factor regulatory networks are specific to microglia clusters

To characterize the regulatory networks of the populations in the dataset, we identified the top transcription factor driven networks (regulons) controlling gene expression in each of the microglia clusters (Figure 3). Each cluster is defined by a specific set of regulons (Figure 3A) supporting the hypothesis that the differential gene expression characterizing each cluster is determined by transcriptional regulation mechanisms. To demonstrate the diversity of regulons predicted to drive gene expression in different clusters we highlight Cluster 3, Cluster 5, Cluster 6, and Cluster 8 (Figure 3B). Each of these clusters shows a different set of regulons that appear as one of the top 10 for that cluster repeatedly across permutations of the analysis. For example, Cluster 5 shares a glycolytic and endolysosomal phenotype with Cluster 6 but does not share the interferon response factor regulons predicted for Cluster 6. In addition, while we observe *IRF1* and *NFKB2* regulons in Cluster 8, the canonical “inflammatory” effector cluster, we see additional (and different) interferon response factor regulons in Cluster 6. The top regulons for other clusters also differ from each other (Figure S6). *MafB*, a transcription factor associated with anti-inflammatory gene regulation, was top of list in Cluster 3^13^. In contrast, regulons directed by transcription factors typically associated with antiviral responses, *IRF7*, and to a lesser extent *IRF3*, were top of the list in Cluster 6 consistent with the observation that these cells are also enriched for nucleic acid recognition and endolysosomal pathways (Figure 3B). Cluster 8 shows significant gene regulation by transcription factors *IRF1* and *NFKB2* (Figure 3B) consistent with the inflammatory profile of this cluster. The top regulons for the APOE ε3/ε3 subset of individuals demonstrate similar unique diversity to those described in the larger dataset again suggesting homology across APOE genotypes (Figure S7). Together, these inferred gene networks and their transcription factor regulons demonstrate the unique diversity of the clusters identified here and provide potential regulatory targets for future studies to investigate.

### Microglia transcriptomic progression takes multiple paths

Experiments in model systems with defined stimuli have demonstrated the potential of microglia to acquire diverse phenotypes. However, understanding the progression and phenotypic switches acquired by human microglia *in vivo* has been challenging. The rich single cell transcriptome data set generated from AD and control cases was employed to investigate how microglia transcriptomic transitions develop by analysis using the Monocle3^37^ trajectory inference method (Figure 4). We asked which cluster may be end state versus transition state as a hypothesis-generating exercise. The resulting branching trajectories suggest that multiple transition states radiate out from HM, the homeostatic marker cluster supporting the premise that “homeostatic” microglia may transition to multiple endpoint phenotypes in humans as predicted by model studies^6,12,13^. We found relationships between clusters that were not immediately apparent when exploring DEGs and GSEA alone. For example, trajectory analysis reveals a branch point where cell progression continues to either the autophagic stress ELN cluster (Cluster 5) or the inflammatory ELN, Cluster 6 (Figure 4). Cluster 5 is adjacent to the senescent cluster (Cluster 9) consistent with the notion that autophagy and senescence are related biological pathways and endpoints. Similar to work by Nguyen *et al*., the motile cluster (Cluster 7) is portrayed as another endpoint^21^.

**Figure 4.**
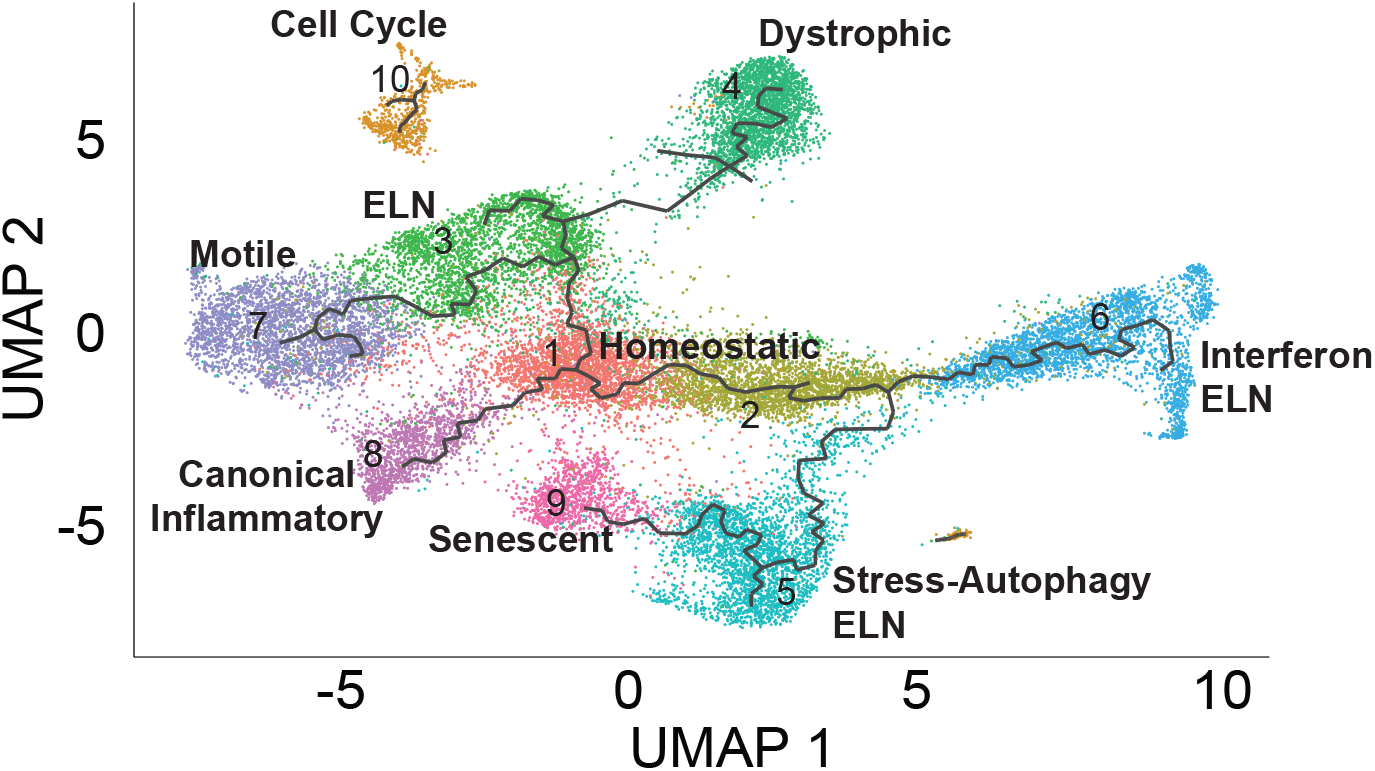
Microglia transcriptomic progression may take multiple paths. Monocle trajectory inference applied to the microglia dataset demonstrates that multiple phenotypic options radiate outward from Cluster 1. Each branch has several potential endpoints suggesting microglia may not progress along a single staged linear trajectory, but instead proceed through one of several transition states to reach various transcriptomic end phenotypes.

### AD cases have an increased proportion of microglia with an inflammatory ELN profile

Both homeostatic marker and canonical inflammatory clusters (HM and Cluster 8 respectively) were equally represented by both AD and control brain. In contrast, we found that Cluster 6 had more AD nuclei than would be expected in our dataset (adjusted *p*=0.006), suggesting AD relevant processes may be represented in the profile of this cluster (Figure 5A). Multiple AD GWAS have identified risk alleles associated with genes expressed in microglia or myeloid cells^8^. Utilizing a list of 46 genes in SNP loci associated with altered AD risk^36,38^ and a list of ELN genes from a GO ELN dataset, we used GSEA to assess enrichment of these gene sets in the 10 identified microglia clusters. We observed more AD risk genes are differentially expressed in Cluster 6 (Figure 5B) compared to all other clusters (adjusted *p*<0.001). Gene expression of *PICALM, SORL1* and *PLCG2* are significantly lower in Cluster 6 compared to the rest of the microglia, while other genes including *APP, APOE*, and *BIN1* were significantly higher in expression (all adjusted *p*<0.001; Figure 5B). Cluster 6 also demonstrated significantly more ELN genes differentially expressed by GSEA than other clusters (data not shown). This finding suggests that AD is associated with an increased proportion of microglia that demonstrate an endolysosomal, glycolytic, and inflammatory gene expression profile. Additionally, these data confirm previously reported findings that the subpopulation of microglia proportionately altered in AD human brain differs from that identified in mouse models^17,39^. We further sought to address whether a microglia phenotype present in healthy aging was reduced in AD brain. We determined that Cluster 10, the cluster differentially expressing cell cycle regulatory genes, is larger in control brain compared to AD brain (Mixed *APOE* adjusted *p*<0.001; *APOE* ε3/ε3 adjusted *p*<0.001; Figure 5A). Cluster 10 is the smallest cluster in our dataset, and therefore deserves replication; however, these data suggest that AD pathology may involve a detectable reduction in a microglia clusters enriched for cell cycle and DNA repair genes.

**Figure 5.**
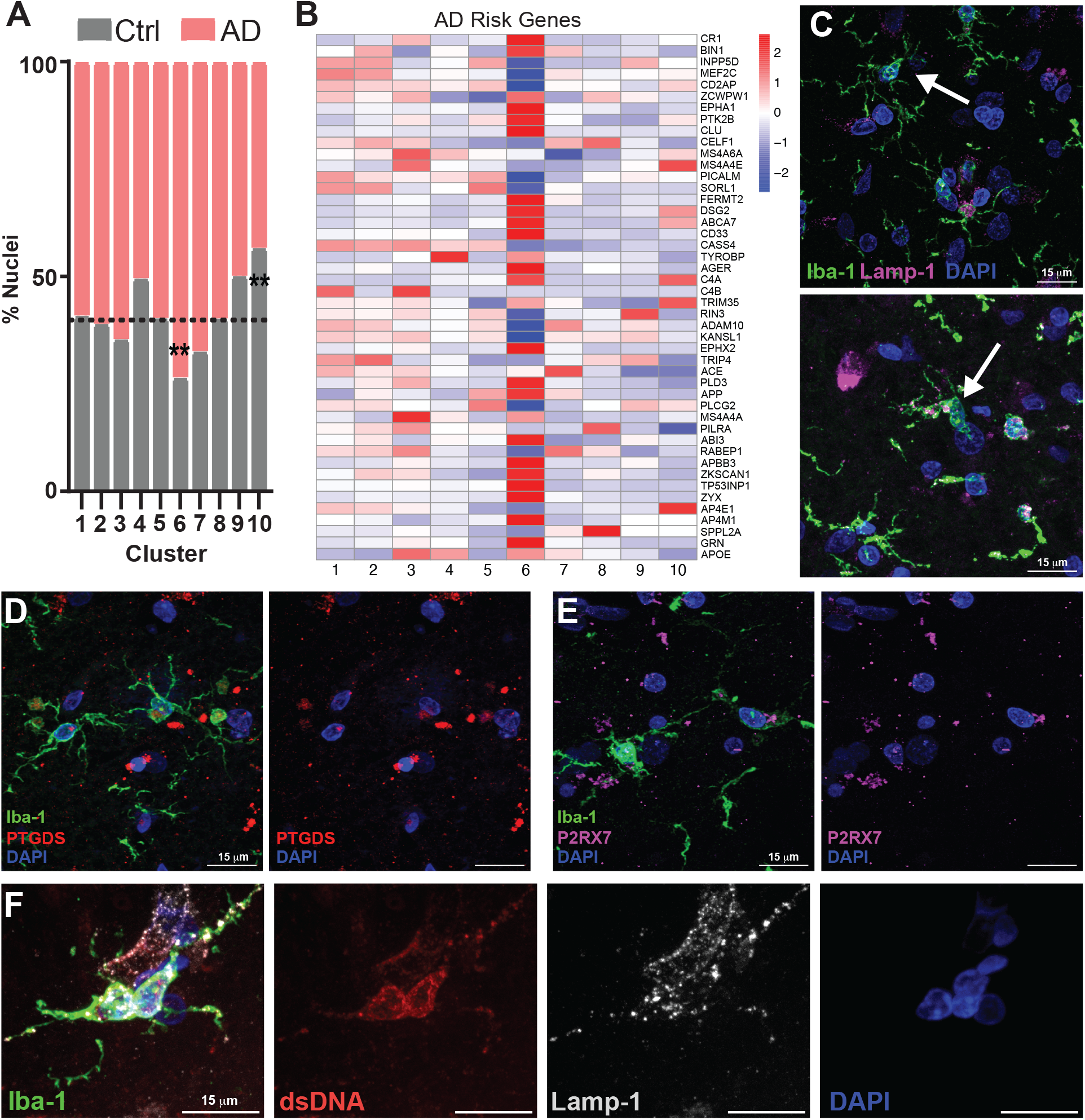
Cluster 6 demonstrates enrichment of AD risk genes and suggests the presence of dysregulated lysosomal and cytosolic DNA regulation in microglia in AD. **A)** Cluster 6 is significantly increased in AD brain, whereas Cluster 10 is increased in control samples. **B)** Heatmap of AD associated risk gene expression across microglia clusters show stronger differential expression in Cluster 6. **C)** Representative images from an AD case demonstrate microglia (green, Iba-1) with heterogeneity in morphology and Lamp-1 signal. Examples of a ramified (top arrow) and greater lysosome (magenta, Lamp-1) signal and an “activated” or less-ramified phenotype (bottom arrow). **D)** Representative microglia (Iba-1 green) with high PTGDS (red) expression a Cluster 6 differentially expressed gene in an AD case. **E)** Representative microglia (Iba-1 green) with high P2RX7 (magenta) expression a Cluster 6 differentially expressed gene in an AD case. **F)** Representative microglia with inreased lysosomal signal (Lamp-1 signal, white) and cytosolic dsDNA (red) in an AD case. **corrected *p*<0.01

### Transcriptomic phenotypes may correspond to microglial morphological phenotype

The transcriptomic data predicts heterogeneity in microglia endolysosomal phenotypes in both aged control and AD brain. Immunostaining for LAMP1, a lysosomal marker, revealed a spectrum of lysosomal phenotypes with varying lysosomal size and number (Figure 5C). Microglia expressing Cluster 6 markers were identified in AD brain by immunolabeling for PTGDS (Figure 5D) and P2RX7 (Figure 5E), both highly expressed genes in that subcluster. Cluster 6 and Cluster 8 microglia both also demonstrate differential expression of genes involved in the detection of DNA/RNA molecules suggests that microglia may be activated by exposure to cytosolic nucleic acids. To assess the presence of cytosolic nucleic acids in microglia, we immunolabeled microglia for double-stranded DNA (dsDNA; Figure 5D). We observed that a subset of microglia with immunoreactivity for dsDNA also contained enlarged lysosomes (Figure S8A,B), while other microglia in the same tissue section had normal lysosome size and no immunoreactivity for dsDNA (Figure S8B). These findings suggest that microglia heterogeneity reflected by gene expression patterns may represent specific morphological phenotypes that can be detected in human tissue.

### An AD specific state exists within the homeostatic marker cluster

As described above, in the HM cluster, AD and aged control microglia were proportionally similar. We reasoned that given the known inflammaging changes associated with human brain molecular profiles, it could be challenging to detect AD specific signatures when studying polarized subpopulations in AD and aged brain tissue. We turned our attention to the complexity within HM. As has been previously reported, we observed that HM is the largest population^17,20,21^. Nevertheless, this population has not been further characterized to understand the impact of AD on gene expression within homeostatic microglia. Our dataset provides depth of coverage and over 50,000 microglia enriched for homeostatic markers, enabling detection of subclusters within the HM microglia population. Sub-clustering HM microglia revealed seven populations with differential gene expression (Figure 6A). We found that in contrast to the larger microglia dataset, there was indeed a subcluster nearly unique to AD cases. This AD specific microglia sub-type, designated Subcluster 1.5 (Figure 6B); was almost exclusively composed of AD microglia, suggesting that it may be uniquely driven by AD pathology (adjusted *p* < 0.001). In contrast, Subcluster 1.4 was overrepresented by control nuclei (Figure 6B; adjusted *p* < 0.05). Gene expression demonstrated very high gene expression of P2RY12 in all HM subclusters including highest expression in Subcluster 1.5 consistent with their location in the HM cluster (Figure 6C). Other gene expression markers of HM Subcluster 1.5 were more specific, including WIPF3, PDE4B, and KCNIP that were identified as differentially expressed genes (Figure 6C). To begin to characterize the putative biological processes represented in these subclusters, we performed GSEA as above. Further, Subcluster 1.5 had a unique profile of enrichment for genes involved in cellular motility and calcium signaling (Figure 6D). Using immunohistochemistry, we validated the presence of double-positive high P2RY12 and PDE4B protein expression in microglia in human AD brain (Figure 6E). We additionally verified these results in our cohort of all APOE ε3/ε3 individuals. We confirmed that again, homeostatic marker microglia could be divided into multiple populations (Figure S9A) with distinct gene expression and that one such cluster was enriched in AD cases (Figure S9B). Together, these data suggest that there exists within the homeostatic marker population a microglia state that is increased in AD brain.

**Figure 6.**
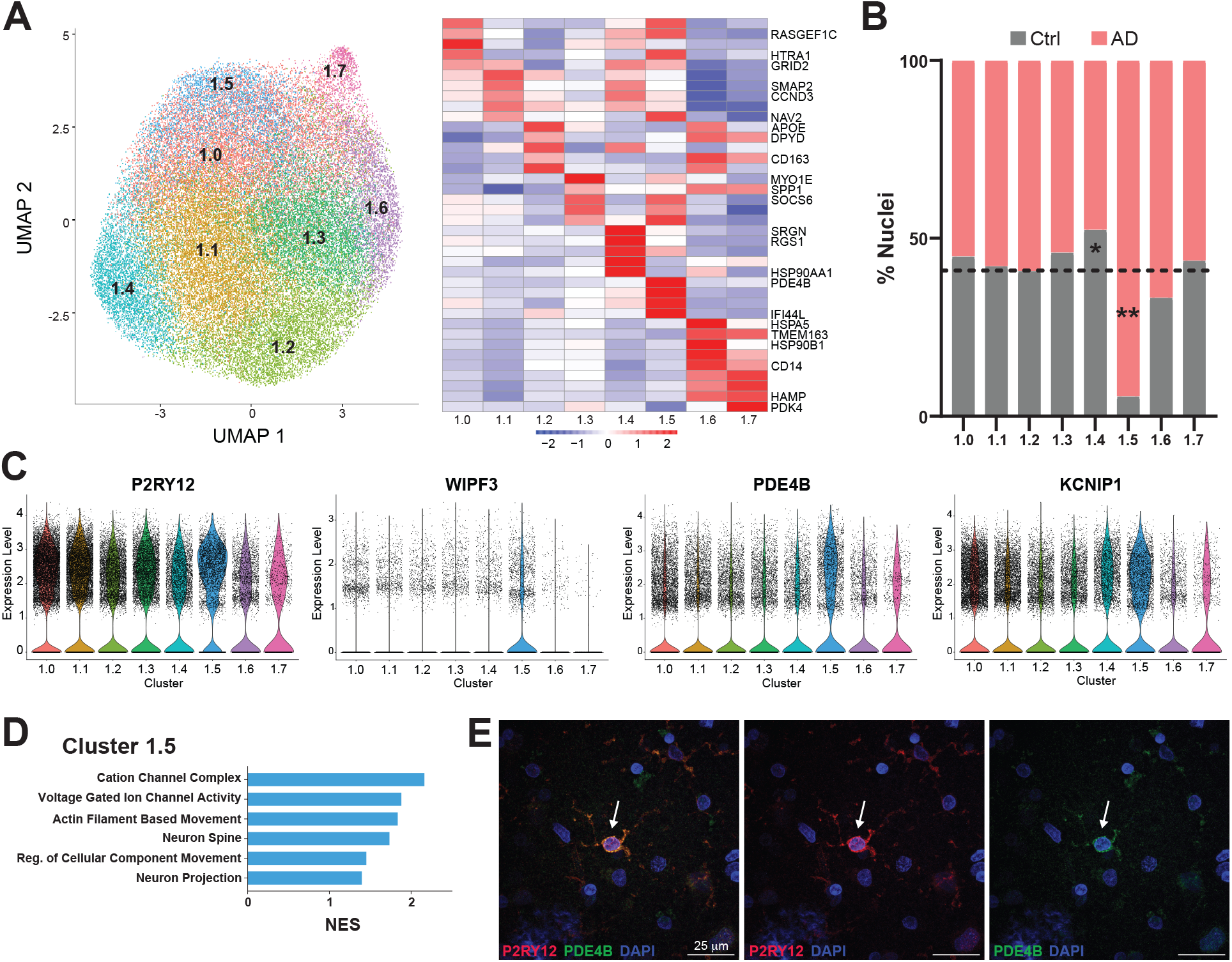
Within microglia expressing homeostatic markers there is a subcluster uniquely enriched in AD brain samples. **A)** Subclustering of Cluster 1 homeostatic microglia revealed 7 subpopulations defined by differential gene expression. **B)** Subcluster 1.5 is greatly expanded in AD brain samples (*p*<0.001), while Cluster 1.4 is increased in control brain samples (*p*<0.05). **C)** Gene expression of P2RY12 demonstrates high expression across the HM subclusters, with highest expression in Subcluster 1.5. Genes differentially expressed in Subcluster 1.5 such as *WIPF3, PDE4B*, and *KCNIP* also demonstrate high expression. **D)** Pathway enrichment in Subcluster 1.5 demonstrates unique enrichment for motility, ion channel activity, and neuron-related processes. **D)** Immunohistochemistry validates the presence of double-positive high P2RY12 and PDE4B expressing microglia in AD brain. NES = Normalized Enrichment Score *corrected *p*<0.05 **corrected *p*<0.01

## Discussion

This study identified ten unique microglia clusters from aged human brain. These included previously described homeostatic, senescent, and inflammatory microglia phenotypes, as well as novel clusters of transcriptional specification. We describe the diversity of microglia clusters with endolysosomal gene signatures one of which is enriched with nucleic acid recognition and interferon regulation genes. Inferred gene networks demonstrated that individual clusters were predicted to be driven by distinct transcription factors lending further support to the putative functional diversity of clusters. Using trajectory inference analysis we observe transitions in microglia phenotypes and predicted relationships that can be tested *in vitro* such as the association of an autophagy enriched cluster with the senescent cluster. Microglia identified as expressing homeostatic markers were themselves transcriptomically diverse. The most differentially present cluster between AD and controls was a homeostatic marker subcluster characterized by calcium activation, response to injury and motility pathways.

Microglia clearance of pathologic proteins including amyloid-beta and tau are established features of AD pathophysiology. The endolysosomal network is critical to maintaining cellular homeostasis and has long been implicated in AD pathogenesis though less is known about the function and dysfunction of ELN components in AD microglia^40^. This study resolved microglia enriched for endolysosomal pathways into three clusters transcriptionally distinguished by predicted biological function. Together, three ELN clusters, Clusters 3, 5 and 6, represent a diversity of endocytosis, vesicle trafficking, endolysosomal and autophagosome pathways defined within the dataset. Impaired microglial endolysosomal function is often discussed in the context of insufficient amyloid clearance^40,41^ though may have additional consequences contributing to AD. The endolysosomal system in myeloid cells also plays a critical role in identifying and processing foreign microbes including initiation of TLR and interferon signaling^25,42^. Cluster 6 was characterized by lysosomal and vesicular function pathway enrichment and concomitantly increased expression of interferon regulatory factor and inflammasome activation genes^32,34,43^. Interferon signaling has been implicated in the response to amyloid fibrils contributing to synapse loss^44^. While there is not a single “DAM” phenotype in human AD, as has been well described in murine neurodegnerative models, there is a type 1 interferon reponse cluster distinct from a dystrophic or canonical inflammatory phenotype^14,15,30,45^. We hypothesize that IRF expression here may reflect exposure to danger associated molecular pattern molecules including nucleic acids which was supported by the presence of microglia with immunoreactivity to cytosolic dsDNA and amoeboid morphology. These findings are in line with murine and *in vitro* studies which show that amyloid fibrils containing nucleic acids can induce a type I interferon response and subsequent synapse loss^44^. Song *et al*. report a key role for the ELN in nucleic acid degradation and the interferon response to DNA damage after release of nuclear DNA into the cytosol^46 43^. These *in vitro* studies underscore the role of the ELN in the resolution of cytosolic nucleic acid responses. Numerous studies report differential expression of AD risk genes, many of which are involved in the endolysosomal system^36^, in brain tissue from both mouse and human AD tissue compared to control^16,17,20,21,30,47,48^. In this study we identified a microglia phenotype, Cluster 6, in which AD risk genes were most strongly differentially regulated and may be contributing to that observed differential expression. By focusing on specific microglial clusters, and thus biological contexts in which risk genes are regulated, we may inform efforts to understand how AD risk genes lead to AD. This is the first report describing a specific cluster of microglia with ELN profile that is both larger in AD brain and enriched for expression of genes associated with both AD risk and interferon activation.

We leveraged the depth of our dataset to apply trajectory methods and infer the potential relationships between microglial transcriptional phenotypes. Dystrophic (Cluster 4) and senescent (Cluster 9) clusters emerge as “end-states” which, while requiring validation in larger datasets, demonstrates that computational analysis such as trajectory inference can map microglial phenotypes consistent with predictions from previous empirical data^6,12,13^. Nevertheless, the nature of trajectory analysis means that these data should be viewed as hypothesis-generating. After observing three microglia clusters with ELN phenotypes, the question arose as to whether they represent three clusters along one lineage. Instead, we found that the autophagic stress and inflammatory ELN clusters (Cluster 5 and Cluster 6 respectively) represented a branch point from homeostatic clusters. Cluster 3 instead appeared to be a transition between homeostatic and alternate transcriptomic endpoints, the motile or dystrophic clusters. These findings are similar to Nguyen *et al*. who also demonstrate a dystrophic cluster as an endpoint preceded by an intermediate state^21^. Our analysis nominates genetic regulators of each cluster and together these data can be used to guide further studies to test the plasticity or reversibility of microglia phenotypes. We have described the transcriptomic heterogeneity as clusters of subpopulations, though in the living brain, microglial plasticity may manifest as a continuum rather than discrete populations^49^. Furthermore, unlike other cell types that are terminally differentiated, it is plausible that microglia may transition in and out of transcriptomic states, underscoring the “snapshot” nature of tissue omics.

It has been more challenging to consistently identify an AD specific microglia cluster in the aged human AD brain than in transgenic animal AD models. This is likely in part due to the overlap in microglial responses to exposure to AD pathology, aging, environmental stressors as well as the diversity of human patients themselves. However, when we analyzed clusters that had not already been driven to a reactive phenotype we then detected an AD specific subpopulation, Subcluster 1.5, in the dataset. The most highly differentially regulated gene in Cluster 1.5 is *PDE4b*, a phosphodiesterase implicated in cognitive function^50^ and myeloid cell inflammatory activation^51–53^. PDE4b regulates levels of cAMP which in turn has been shown to modulate microglia surveillance behavior^54^. The enrichment of motility and calcium signaling pathways by GSEA may indicate responses to neuronal activity and extension of microglial processes active in the AD brain^49^. Furthermore, the elevated levels of *P2RY12* would be consistent with a phenotype of early response to responding to damage signals and extending processes to injured neurons^55–57^. Our data underscores the heterogeneity of microglia expressing canonical “homeostatic” markers or included in a “homeostatic” cluster. Differentially expressed genes and pathways enriched in the AD specific Subcluster 1.5 may offer clues into the early or subtle microglial phenotype changes in response to pathologic protein, prior to acquiring a more immunologically activated phenotype.

As expected, given its frequency and strong association with AD, we observe more copies of the *APOE* ε4 allele in AD cases than controls within our sample reflecting the availability of tissues from our brain bank. *APOE* ε4 is a key component to AD pathogenesis in many patients. Therefore, the degree of similarity between microglia subpopulations when comparing the mixed *APOE* group and the *APOE* ε3/ε3 cohort was notable. However, the limited sample size remains a caveat to drawing strong conclusions.

Like all snRNAseq studies, there are limitations to our study. While PU.1 sorting provides a way to increase microglia nuclei for greater depth of resolution at an individual level, it potentially selects for a specific population of microglia. DEGs identified utilizing clusters defined by the same data are not necessarily properly controlled^58^, however our large dataset will allow others to utilize our clusters to mitigate this in the future. Although gene expression is a useful molecular tool for cellular subtyping, it does not always directly describe protein expression, localization, or activity^59^. Future studies assaying for a panel of proteins based upon the transcriptomic signatures reported here will be valuable for both validation as well as the spatial correlation with pathological features. Assessment of microglial phenotypes across brain regions can provide further context to our understanding of phenotypic heterogeneity. Another significant limitation is the use of autopsy brain tissue. It is possible that events just prior to death or post-mortem changes contribute to the expression changes measured here. To mitigate the variability of tissue quality, we have only selected tissues with a pH greater than 6.0 and PMI less than ten hours. Nevertheless, this novel work demonstrates the improved resolution that can be achieved in autopsy brain tissue revealing phenotypes recognized *in vitro* and in other inflammatory models, but previously unidentified in human AD brain.

In this study we identified and assessed microglia states from isolated human post-mortem brain microglial nuclei. Genes suggestive of an “aging signature” are expressed in all clusters in this study^18^ consistent with our older age cohort. Inflammaging^60^ may not only confound interpretations of gene expression profiles attributable to AD, but may also contribute to the disease mechanisms hypothesized to drive AD. Additional studies exploring differences between younger controls and early-onset AD may also help to explore the aging, inflammaging, and AD specific signatures. These data identify candidate genes and pathways driving microglia responses to AD across a spectrum of microglial activation phenotypes associated with specific transcription factor regulons. Our identification of multiple internalization and trafficking clusters with varying metabolic and inflammatory phenotypes provides a platform for further studies to replicate and investigate. Finding a homeostatic marker expressing subcluster that is unique to AD, while requiring replication, also suggests that AD changes and thus molecular pathways driving AD can be identified in cells that have not yet been fully explored. The cluster-specific alterations in composition, gene expression, and gene regulation in AD brain provide additional information to support tailored targeting of microglial physiological responses that will be critical moving forward in neurodegenerative disease therapeutics.

## Methods

### Human Brain Tissue

Dorsolateral prefrontal cortex (dlPFC) tissue from human brains was obtained from the Neuropathology Core of the Alzheimer’s Disease Research Center (ADRC) at the University Washington (UW) following informed consent approved by the UW Institutional Review Board (IRB). Patients (n=12) were confirmed post-mortem to have AD pathology (ADNC score of 2-3; Table 1). Control individuals (n=10) had low or no neuropathology post-mortem (ADNC score 0-1; Table 1).

**Table 1.**
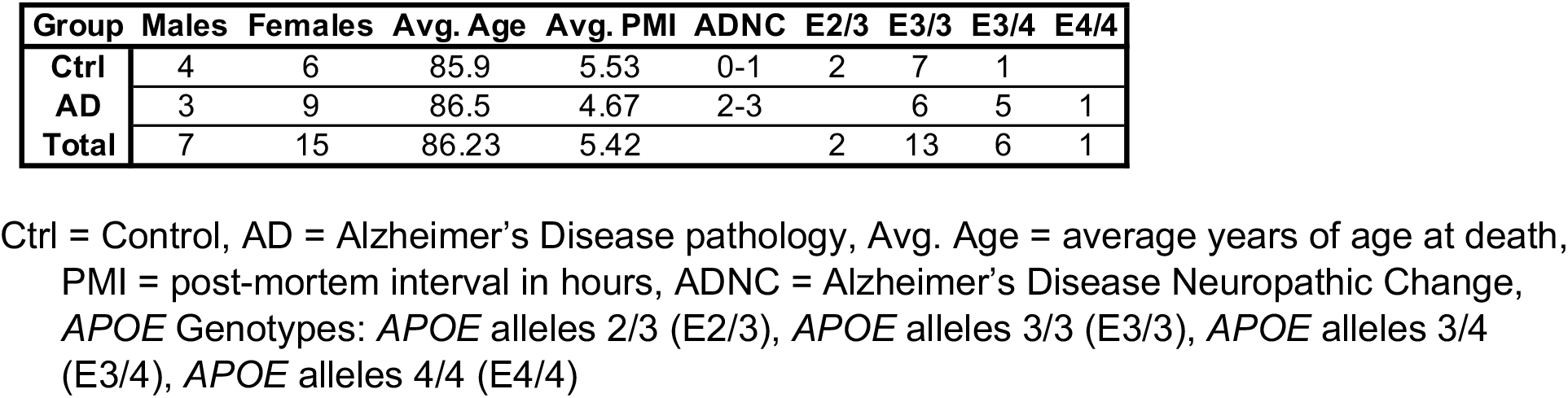
Post-mortem brain sample demographics.

Brain samples were flash-frozen and stored at -80ºC. Criteria for inclusion included post-mortem interval (PMI) ≤ 10hrs, low comorbid pathology (Lewy Bodies and hippocampal sclerosis), and a brain pH at autopsy ≥ six.

### Isolation of Nuclei for Unsorted snRNA-seq

Nuclei from brain samples were isolated using protocols adapted from 10X Genomics Demonstrated Protocols and De Groot *et al*.^61^. Briefly, four 2mm punches of dlPFC gray matter were collected using a biopsy punch (Fisher Scientific, Waltham, MA) into a 1.5mL microcentrifuge tube on dry ice. All buffer recipes and reagents can be found in Table S4. Nuclei isolation used nuclei lysis buffer (NLB). The nuclei in nuclei suspension solution (NSS) was layered onto 900μL of Percoll/Myelin Gradient Buffer (PMGB)^61^. The gradient was centrifuged at 950xg for 20min at 4°C with slow acceleration and no brake. Myelin and supernatant were aspirated and the nuclei pellet was resuspended in Resuspension Buffer (RB) at a concentration of 1000 nuclei/μL and proceeded immediately to single-nuclei RNA sequencing (snRNA-seq).

### Isolation of Nuclei for Fluorescence-activated Nuclei Sorting (FANS)

Briefly, 200-250mg of dlPFC was collected into a 1.5ml microcentrifuge tube on dry ice. Brain tissue was homogenized as above. The homogenate was incubated at 4ºC under gentle agitation for 10min, pelleted at 500xg for 7min at 4ºC and resuspended in 900μL PMGB supplemented with protease and phosphatase inhibitors. The suspension was gently overlaid with 300μL NSS supplemented with protease and phosphatase inhibitors. The gradient was centrifuged at 950xg for 20min at 4ºC with slow acceleration and no brake. The myelin and supernatant were aspirated and the nuclei pellet proceeded immediately to FANS.

### Fluorescence Activated Nuclei Sorting (FANS)

Nuclei were washed with cold FANS media (10% fetal bovine serum (FBS), 10mM HEPES, 100μM ATA, 10% 10X HBSS, 0.5% Protector RNase Inhibitor, protease and phosphatase inhibitors, and 1% saponin in nuclease-free water) and resuspended in FANS media at a concentration of 2-2.5×10^6^ nuclei/mL. Nuclei were incubated with 1% Human Fc Block (clone Fc1.3216, BD Biosciences, San Jose, CA) on ice for 10min. Nuclei were labeled with either anti-PU.1-PE (clone 9G7, 1:50, Cell Signaling Technology, Danvers, MA) or IgG-PE isotype control (clone DA1E, 1:50, Cell Signaling Technology) for 4hours on ice followed by three washes with cold FANS media and resuspended in FANS media supplemented with 10μg/mL DAPI (Sigma-Aldrich). Nuclei were sorted using a FACSAria III (BD Biosciences) until 30,000 PU.1-positive nuclei were collected. Sorted nuclei were centrifuged at 1,000xg for 10min at 4ºC. The nuclei pellet was resuspended in RB at a concentration of 1000 nuclei/μL and proceeded immediately to snRNA-seq.

### Single Nuclei RNA-Sequencing (snRNA-seq)

Single nuclei libraries were generated using the Chromium Next GEM Single Cell 3′ GEM, Library and Gel Bead Kit v3 (10x Genomics, Pleasanton, CA) according to the manufacturer’s protocol and a target capture of 10,000 nuclei. Gene expression libraries were sequenced on the NovaSeq 6000 platform (Illumina, San Diego, CA).

### Alignment and Quality Control

Gene counts were obtained by aligning reads to the hg38 genome (GRCh38-1.2.0) using CellRanger 3.0.2 software (10x Genomics). Reads mapping to precursor mRNA were included to account for unspliced nuclear transcripts. The majority of our analysis was performed in R^62^. Droplets from 22 PU.1 sorted samples were combined using Seurat v3.3^63^. Unsorted and PU.1 sorted droplets isolated from the same 4 subjects were combined using Seurat and analyzed in the same manner. Droplets containing less than 350 UMIs, less than 350 genes, or greater than 1% mitochondrial genes were excluded from analysis. Ambient RNA was removed from the remaining droplets using SoupX^64^. Droplets containing multiple nuclei were scored using Scrublet^65^, and removed. 200,948 nuclei with an average of 1,787 genes per nucleus remained in the dataset for further analysis.

### Normalization and Cell Clustering

Normalization and clustering of the nuclei were performed using Seurat v3.3^63^. Data were normalized for read depth and mitochondrial gene content was regressed out using Seurat’s SCTransform^66^. Individual sample variability was removed using Seurat’s Anchoring and Integration functions^63^. The top 5000 variable genes were kept. 15 principal components (PCs) were used to create a shared nearest neighbors graph with k=20. The modularity function was optimized using a resolution of 0.2 to determine clusters using the Louvain algorithm with multilevel refinement to determine broad cell-types. Each cluster met a minimum threshold of 30 defining DEGs and was comprised of nuclei from >10% of the cohort (more than two individuals).

Clusters were annotated for cell-type using manual evaluation for a set of known genetic markers^67^. A new Seurat object was made containing only the microglia nuclei (N = 127,371). Normalization, individual variability removal, integration, and shared nearest neighbors graph creation were repeated as above on the microglia nuclei. 20 PCs were chosen to account for a significant amount of the variance. Clusters were determined using the Leiden algorithm^68^ with `method=igraph` and `weights=true`. Clusters were highly conserved across analysis by Louvain, Louvain with multilevel refinement, and Leiden algorithms. HM subclustering for both the 22 sample dataset, and the APOE !3/!3 allele dataset occurred after normalization, individual variability removal and integration as before and was performed using 10 PCs and the Louvain with multilevel refinement algorithm. Distribution of nuclei within each cluster was calculated using the ‘`chisq.test`’ function in R^62^ to compare the actual percentage of nuclei from either the AD or control group within the cluster to the expected proportion of nuclei that would be contributed based on dataset composition. P-values from the chi-squared tests were adjusted using FDR and considered significant if adjusted *p*<0.05.

### snRNA-seq Differential Gene Expression and Gene Set Enrichment Analyses

Differential gene expression analysis of the clusters was performed with the MAST algorithm. Genes tested had expression in at least 25% of the nuclei in the cluster. Differentially expressed genes (DEGs) had an FDR adjusted p-value<0.05 and a log fold change>1.25. Cluster 1 was annotated as inactivated, often referred to as “homeostatic” in single nuclei studies of microglia. Differential gene expression analysis was repeated as above comparing each other cluster to Cluster 1. Gene set enrichment analysis (GSEA) was performed in ClusterProfiler^69^ modified to use a set seed for reproducibility, using the GO, KEGG and Biocarta pathway sets, version 7.2. Enriched pathways had an FDR adjusted *p*-value<0.05. We considered pathways to be representative if significant results included similar genes and biological functions in at least two of the three major databases (GO, KEGG, Reactome).

### Gene Ontology Biological Process Clustering

A complimentary approach to gene set enrichment analysis (GSEA) is to perform biological process ontology clustering to identify a more extensive set of terms associated with the gene list. To further characterize the ELN clusters (3, 5, and 6) we implemented this approach to get a more refined examination of the biologically linked process driven by each cluster. We employed several different approaches to perform this analysis, ultimately choosing the Cytoscape network clustering application ClueGO^70,71^, which was employed extensively in recent years for this purpose^72–75^. The advantage of this network approach is that GO term enrichment is selected based upon networks derived from kappa-score between terms based on similarity of genetic contribution to the particular term. This creates a network of genetically linked processes that goes beyond a singular GO term hit, providing greater confidence that a basic biological process is being impacted based on the differential expression of a particular set of genes. Clusters 3, 5 and 6 were each independently submitted for analysis, using a kappa-threshold of 0.4 to optimize the biological process connections. Each term drawn into the network was initially filtered for multiple testing corrections threshold of *p*<0.05, and hierarchically weighted for terms with a Benjamini-Hochberg correction value of *p*<0.01. As this procedure is performed for validation and visualization, we trimmed networks of lesser ranked terms to permit ease of visualization.

### Trajectory and Lineage Analysis

Trajectory analysis was performed using Monocle3^37^ on multiple permutations of our downsampled dataset. The data were downsampled to 1000, 2000, 3000, or 5000 nuclei per cluster, transferred to a cell dataset (cds) object and Monocle3 “`learn_graph`” was run. PCA and UMAP embeddings were extracted from the Seurat object. We applied the algorithm both with and without a defined starting point. The 5000 nuclei per cluster downsampled data began to break down the ability of Monocle to form a consistent trajectory, where the 1000, 2000, and 3000 multiple permutations consistently formed something similar to the representative image in Figure 5 (3000 nuclei per cluster).

### Gene Regulatory Network Inference

Regulons were inferred using the SCENIC workflow in python (pySCENIC)^76,77^. We randomly selected 2000, 3000, or 5000 nuclei in each cluster or if they have less than this number all the nuclei in the cluster to reduce the computational time and have proportional representation of all the clusters. We made 5 subsets of each combination and repeated the analysis twice for each subset to assess the consistency of the regulons in the analysis. First, we used normalized counts with highly variable genes to generate the co-expressing regulatory network modules using the machine learning algorithm GRNBoost2 with function “`grn`” and default settings^78^. Second, the modules were filtered using the “`-ctx`” function, which uses cis-regulatory motif analysis {RcisTarget} to keep only modules enriched for putative target genes of the transcription factor. Regulons are identified by combining multiple modules for a single transcription factor. Third, the AUCell function was used to calculate the regulon activity for each nucleus. Regulon specificity scores were calculated for each regulon in every cluster. Ranking specificity scores identified the top 10 regulons for a specific cluster for a given subset of the dataset, and the consistency of the findings across subsets.

### Immunostaining of Human Tissue

Dissected tissue from the dorsolateral prefrontal cortex of the 22 cases in the cohort were fixed with paraformaldehyde and paraffin embedded. Samples were sectioned at 15um and deparaffinized prior to immunostaining. Slides were boiled in citrate buffer (Sigma CAT#C9999) for 20 minutes then transferred into blocking buffer (10% donkey serum, 0.1% Trition X-100, and 0.05% Tween-20 in TBS) for one hour at room temperature. Slides were incubated in primary antibodies (Anti-LAMP1 1:100 Invitrogen CAT#14-1079-80; anti-Iba-1 1:250 Abcam CAT#ab5076; anti-dsDNA 1:250 Millipore CAT#MAB1293; anti-PTDGS/PGD2 R&D Systems CAT#MAB10099 1:100; anti-P2RX7 Santa Cruz CAT#sc-514962 1:100; anti-P2RY12 Alomone CAT#APR-012 1:50; anti-PDE4B LSBio CAT#LS-C173292-100 1:50) overnight at 4ºC. Slides were rinsed three times in TBS-T for five minutes prior to secondary antibody (Thermofisher Alexa Fluor 488 Donkey anti-Goat CAT# A11055; Thermofisher Alexa Fluor 555 Donkey anti-Mouse CAT#A31570; Alexa Fluor 555 Donkey anti-Rabbit 555 CAT#A31572; Alexa Fluor 647 Donkey anti-Mouse CAT#A31571; or Alexa Fluor 647 Donkey anti-Rabbit CAT#A32795) incubation for one hour at room temperature. Slides were then stained with DAPI (1:1000 Millipore CAT#D8417) for five minutes followed by three, five minute, TBS-T washes. True Black (Fisher Scientific CAT#NC1125051) diluted 1:20 in 70% ethanol was added to the slides for 50 seconds, followed by two additional five minute washes in TBS prior to being mounted with Fluoromount-G (Southern Biotech CAT#0100-01). Slides were imaged using either an Olympus Fluoview-1000 confocal microscope, or a spinning disk confocal microscope (Nikon A1R with Yokogawa W1 spinning disk head) with 40X and 100X oil objectives, and the maximum projection of z-stack images were generated. Images were despeckled using Fiji.

## Acknowledgements

Funding for these studies was provided by: RF1AG063540 and P30AG066509. Additional support was provided by the Ellison Foundation. KEP was supported by NIA 5T32-AG052354-02. Autopsy materials used in this study were obtained from the University of Washington Neuropathology Core, which is supported by the Alzheimer’s Disease Research Center (AG05136) and the Adult Changes in Thought Study (AG006781). This work was facilitated by the use of the advanced computational, storage, and networking infrastructure provided by the Hyak supercomputer system at the University of Washington. Conceived and designed experiments: SJ, KJG, KEP, JEY, CLS, BL, GAG. Performed the experiments: KJG, CLS, AS, SR, CDK. Analyzed data: KEP, KJG, SM, AS, KLC, JW. Biostatistics oversight: WS, AShojaie. Wrote the manuscript: KEP, KJG, SJ, SM, AS. Edited the manuscript: CLS, WS, AShojaie, JY, EEB, GAG. Provided code: WS, SM, KLC, NSM, RYK, LH. Conceptualized the research and collaborations: SJ, GAG, JEY, EEB, BL.

## Code availability

The scripts used to generate our analyses are available at (https://github.com/keprater/jayadevlab_pu.1_project). The container needed to run the scripts with the same Seurat and other package versions as are used in the code is available for download here: https://hub.docker.com/layers/jayadevlab/keprater/jayadevlab/6.3/images/sha256-d9b8f6fb69261fbdd45b09b2603c262497d00f09e7310951024a42761b8a6670?context=explore.

## Data availability

The data are available from Synapse.

