## Supplemental Figures and Tables for "Transcriptomically unique endolysosomal and homeostatic microglia populations in Alzheimer’s disease and aged human brain"

### Prater, Green, et al. Supplemental Tables/Figures

Supplemental Table 1: Complete demographic information on the cohort

| Sample | Study Designation | Coded AGE | SEX | Race | APOE Genotype | PMI | ADNC Score |
|---|---|---|---|---|---|---|---|
| 1 | AD | 90+ | F | White | 3,3 | 5.08 | 3 |
| 2 | AD | 74 | F | White | 3,4 | 4.42 | 3 |
| 3 | AD | 90+ | F | Mixed | 3,3 | 5.08 | 2 |
| 4 | AD | 90+ | F | White | 3,3 | 3.75 | 3 |
| 5 | AD | 60 | F | Unknown/Unreported | 3,3 | 5.75 | 3 |
| 6 | AD | 86 | F | White | 3,3 | 8.07 | 2 |
| 7 | AD | 87 | F | White | 3,4 | 4.87 | 3 |
| 8 | AD | 90+ | F | White | 3,4 | 4.25 | 3 |
| 9 | AD | 77 | F | White | 4,4 | 3.33 | 3 |
| 10 | AD | | M | White | 3,3 | 3.33 | 3 |
| 11 | AD | 83 | M | White | 3,4 | 3 | 3 |
| 12 | AD | 90+ | M | White | 3,4 | 4.58 | 3 |
| 13 | Ctrl | 90+ | F | Hispanic / Latino | 2,3 | 4.33 | 0 |
| 14 | Ctrl | 90+ | F | White | 3,3 | 3.75 | 1 |
| 15 | Ctrl | 90+ | F | White | 3,3 | 6.97 | 0 |
| 16 | Ctrl | 80 | F | White | 3,3 | 8.13 | 0 |
| 17 | Ctrl | 90+ | F | White | 3,3 | 7.72 | 1 |
| 18 | Ctrl | 70 | F | White | 3,4 | 7 | 1 |
| 19 | Ctrl | 74 | M | White | 2,3 | 4.83 | 0 |
| 20 | Ctrl | 84 | M | White | 3,3 | 3.92 | 1 |
| 21 | Ctrl | 90+ | M | White | 3,3 | 8.17 | 1 |
| 22 | Ctrl | 82 | M | White | 3,3 | 7.75 | 1 |

Ctrl = Control, AD = Alzheimer's Disease pathology, Coded Age = age at death in years. Ages greater than 90 are coded 90+ to maintain anonymity, F = Female, M = Male, Race = self-reported race, *APOE* Genotypes: *APOE* alleles ε2/ε3 (2/3), *APOE* alleles ε3/ε3 (3/3), *APOE* alleles ε3/ε4 (3/4), or *APOE* alleles ε4/ε4 (4/4), PMI = post-mortem interval in hours, ADNC = Alzheimer's Disease Neuropathic Change

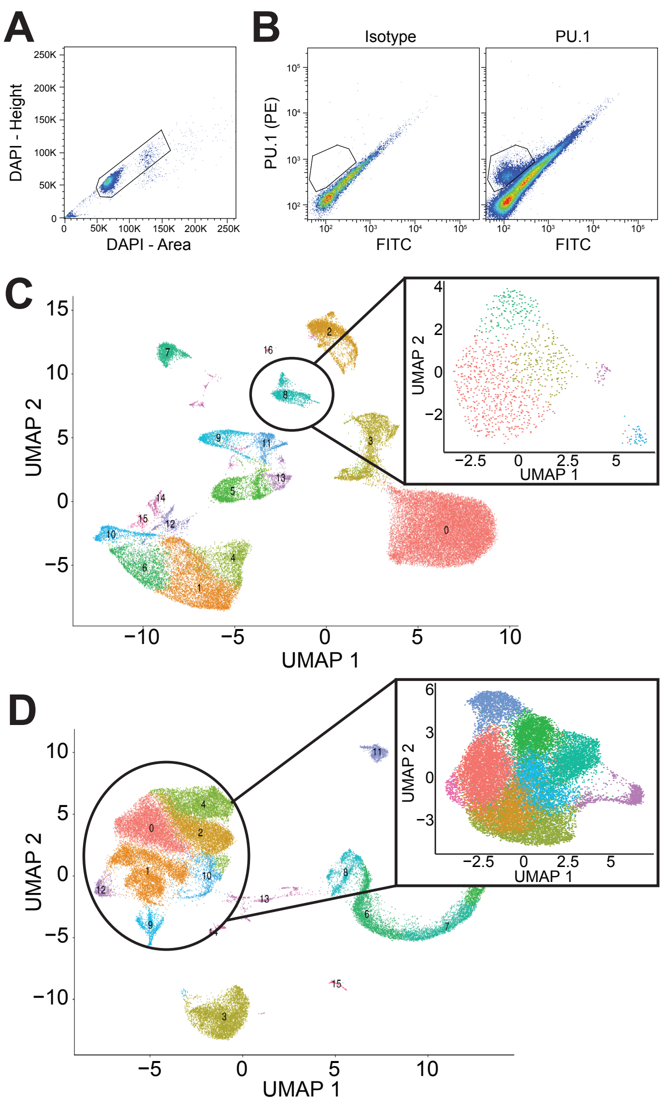


**Supplemental Figure 1. PU.1 enrichment increases the number of microglia nuclei and enhances microglia cluster resolution in snRNAseq studies. A)** Fluorescence-activated nuclei sorting plot of nuclei isolated from dorsolateral prefrontal cortex grey matter of postmortem human brain tissue demonstrate a DAPI-positive population from which later populations are drawn. **B)** Isotype and PU.1 staining examples demonstrating the PU.1 positive population. **C)** Unsorted snRNAseq data from four samples demonstrates multiple brain cell types and a small population of microglia (n = 1032 cells) that can be further subdivided into five clusters. **D)** After PU.1 enrichment, a snRNAseq dataset from the same four individuals contains a larger number of microglia (n = 23,310 cells), and these microglia can be further discriminated into nine clusters.

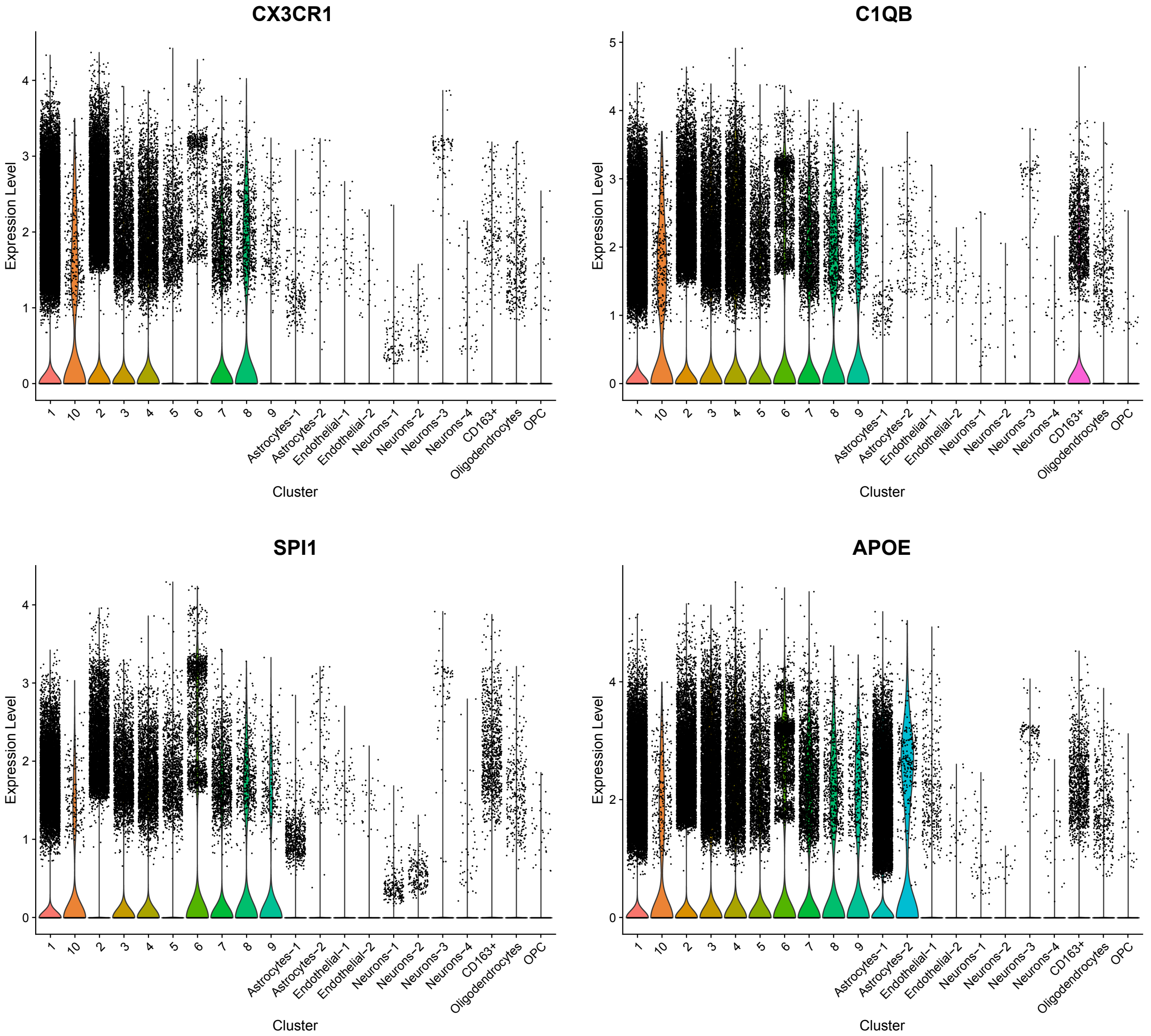


**Supplemental Figure 2. Gene expression of microglia marker genes in the 10 identified microglia subclusters.** Microglia marker genes, CX3CR1, C1Qb, SPI1 (PU.1), and APOE all demonstrate higher expression in the 10 microglia subclusters (numbered 1-10 on the left) than in other cell type clusters (labeled by cell type on the right).

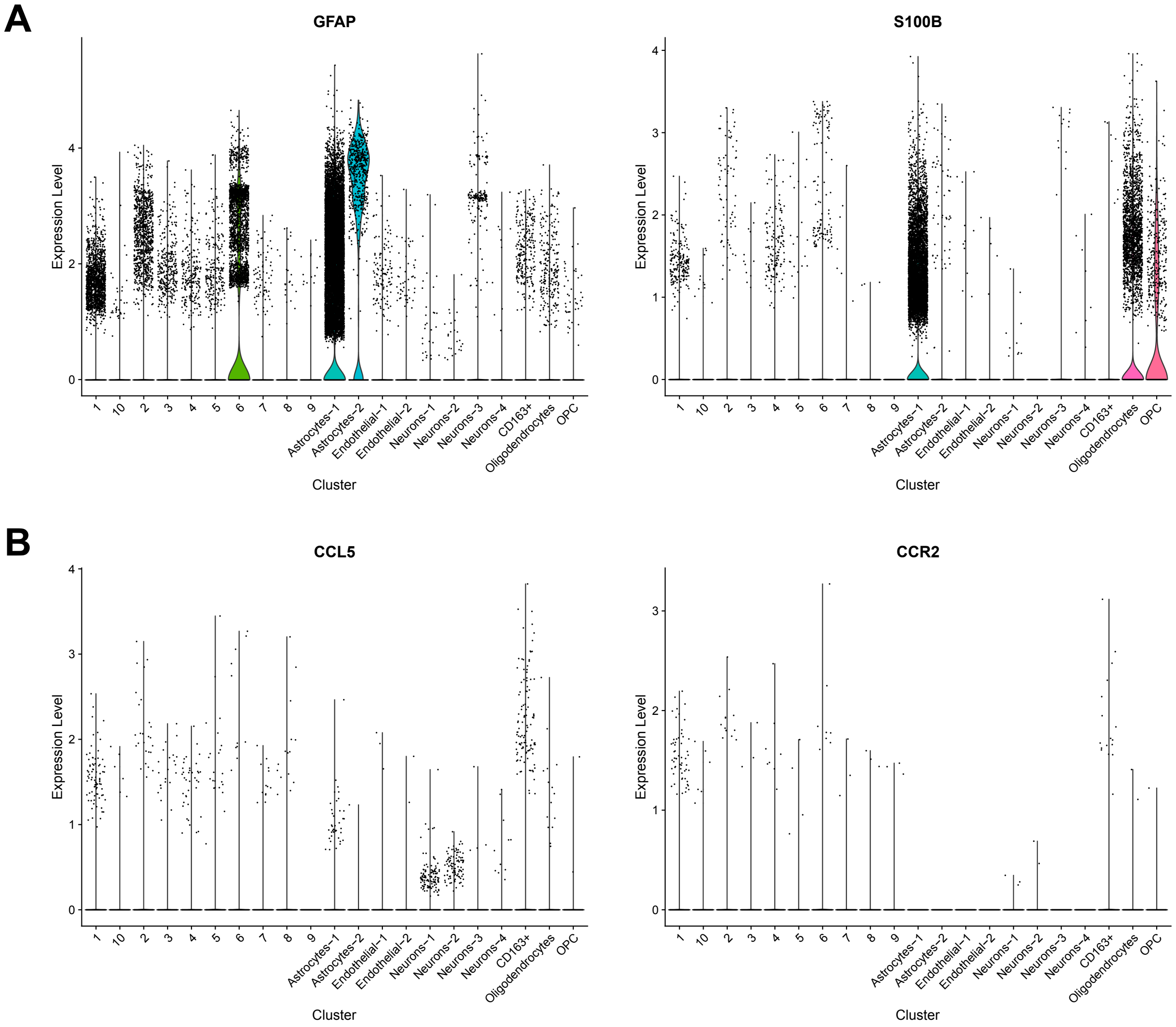


**Supplemental Figure 3. Gene expression of astrocyte and peripheral monocyte marker genes in the 10 identified microglia subclusters. A)** Astrocyte marker genes GFAP and S100B demonstrate higher expression in the two Astrocytes-1 and Astrocytes-2 subclusters than in the 10 defined microglia subclusters. While microglia cluster 6 does have expression of GFAP, it does not have expression of S100B. **B)** Peripheral monocyte markers CCL5 and CCR2 have low expression in our dataset but are most highly expressed by the CD163+ cluster which was not included in our microglia dataset.

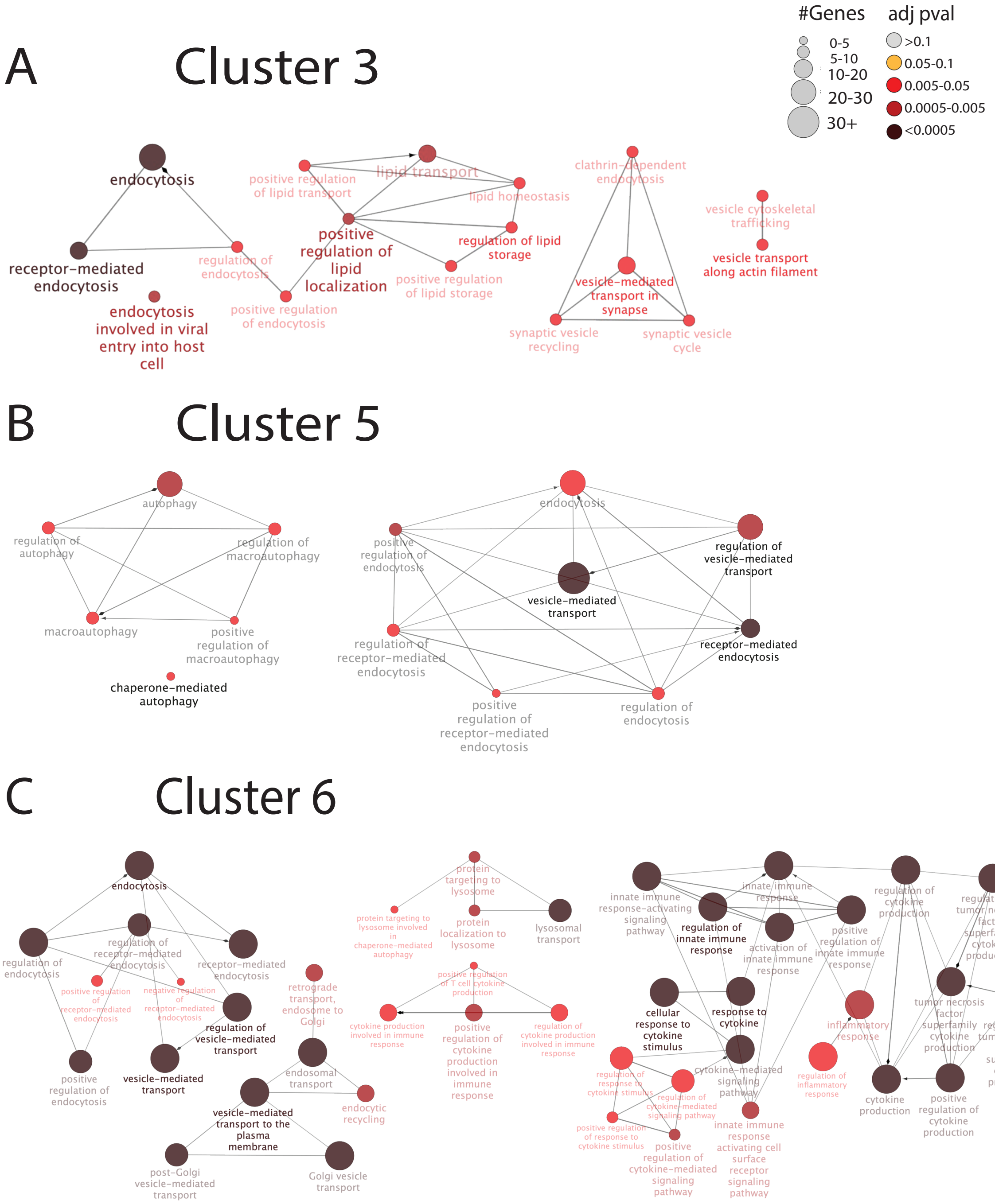


**Supplemental Figure 4. Gene Ontology Network Enrichment of the Endolysosomal Microglia Clusters 3, 5, and 6.** The genes differentially expressed in Clusters 3, 5, and 6 were employed to drive a network-based gene ontology enrichment using the Cytoscape application ClueGO. The nodes are represented within each network based upon two factors: number of genes (size of circle) and statistical significance (bright red = corrected *p*<0.05; dark brown = Benjamini-Hochberg corrected *p*<0.0005). **A)** In Cluster 3, three small subclusters were identified associated with endocytosis, receptor mediated endocytosis, and lipid binding and synthesis (far left); a second subcluster centers on synaptic endocytosis (middle group); and a third involves vesicle transport (far right). **B)** In Cluster 5, a subcluster of terms was identified spanning autophagy, receptor mediated autophagy, and the regulation of these two terms (far left); a second larger cluster identifies endocytosis and receptor-mediated endocytosis, processes linked with autophagic regulation, as well as vesicular transport. **C)** In Cluster 6, three subnetworks were identified within the ELN space, demonstrating active endocytosis, lysosomal processes and transfer between these compartments and the trans-golgi network (left). A second large cluster of terms was identified associated with innate immune function, activation, and regulation, as well as linked inflammatory processes (right side).

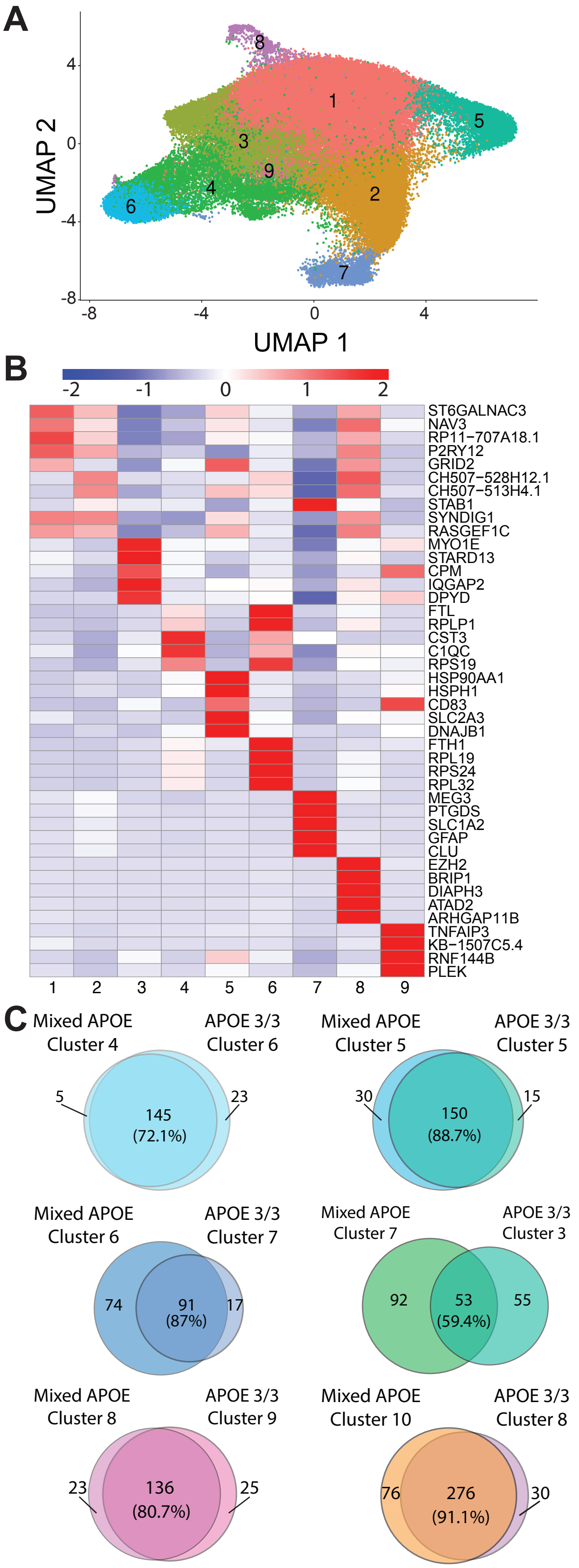


**Supplemental Figure 5. APOE ε3/ε3 genotype does not substantially alter microglial clustering in human autopsy brain. A)** UMAP of unbiased clustering on 13 samples of only APOE ε3/ε3 individuals shows 9 clusters. **B)** Similar to the clusters identified in the mixed *APOE* genotype dataset, the clusters identified in the *APOE* ε3/ε3 genotype dataset are distinct by gene expression. The top 5 genes are displayed for each cluster. **C)** Venn diagrams demonstrating overlap between clusters from the Mixed *APOE* and *APOE* ε3/ε3 cohorts demonstrating significant overlap in gene expression profiles.

| Cluster 4 | % replicates in top 10 | Cluster 7 | % replicates in top 10 | Cluster 9 | % replicates in top 10 | Cluster 10 | % replicates in top 10 |
|---|---|---|---|---|---|---|---|
| JUNB | 96.30 | NFATC1 | 96.30 | PRDM1 | 100.00 | EZH2 | 100.00 |
| STAT1 | 88.89 | DDIT3 | 96.30 | CEBPD | 96.30 | BRCA1 | 96.30 |
| ZNF768 | 85.19 | HIVEP1 | 92.59 | FOXO1 | 81.48 | MAZ | 85.19 |
| E2F6 | 81.48 | MAFB | 85.19 | FOXN2 | 81.48 | MYBL1 | 81.48 |
| PPARA | 77.78 | ATF3 | 77.78 | FOS | 59.26 | FOXN2 | 40.74 |
| JUND | 59.26 | NFIL3 | 74.07 | XBP1 | 48.15 | THAP1 | 40.74 |
| NFE2L1 | 44.44 | ATF5 | 59.26 | PPARD | 48.15 | ZFP64 | 40.74 |
| GABPA | 33.33 | NR3C1 | 55.56 | FOXP2 | 37.04 | ZNF16 | 37.04 |
| KLF9 | 33.33 | E2F3 | 51.85 | XRCC4 | 37.04 | SREBF2 | 37.04 |
| TRIM69 | 33.33 | ZNF235 | 37.04 | RUNX3 | 29.63 | ZBTB2 | 33.33 |

**Supplemental Figure 6. Top transcription factors for each cluster from the full cohort microglia subclusters are unique and represent biological function switches.** These are the transcription factors that most often drove gene expression in Clusters 4, 7, 9, and 10. Values denote the percentage of replicates of permutations of the dataset where that transcription factor was unique to the given cluster. Note that while a few similar transcription factors are seen in multiple clusters, particularly the most prevalent transcription factors are unique, and are representative of gene expression driving biological functions in these clusters.

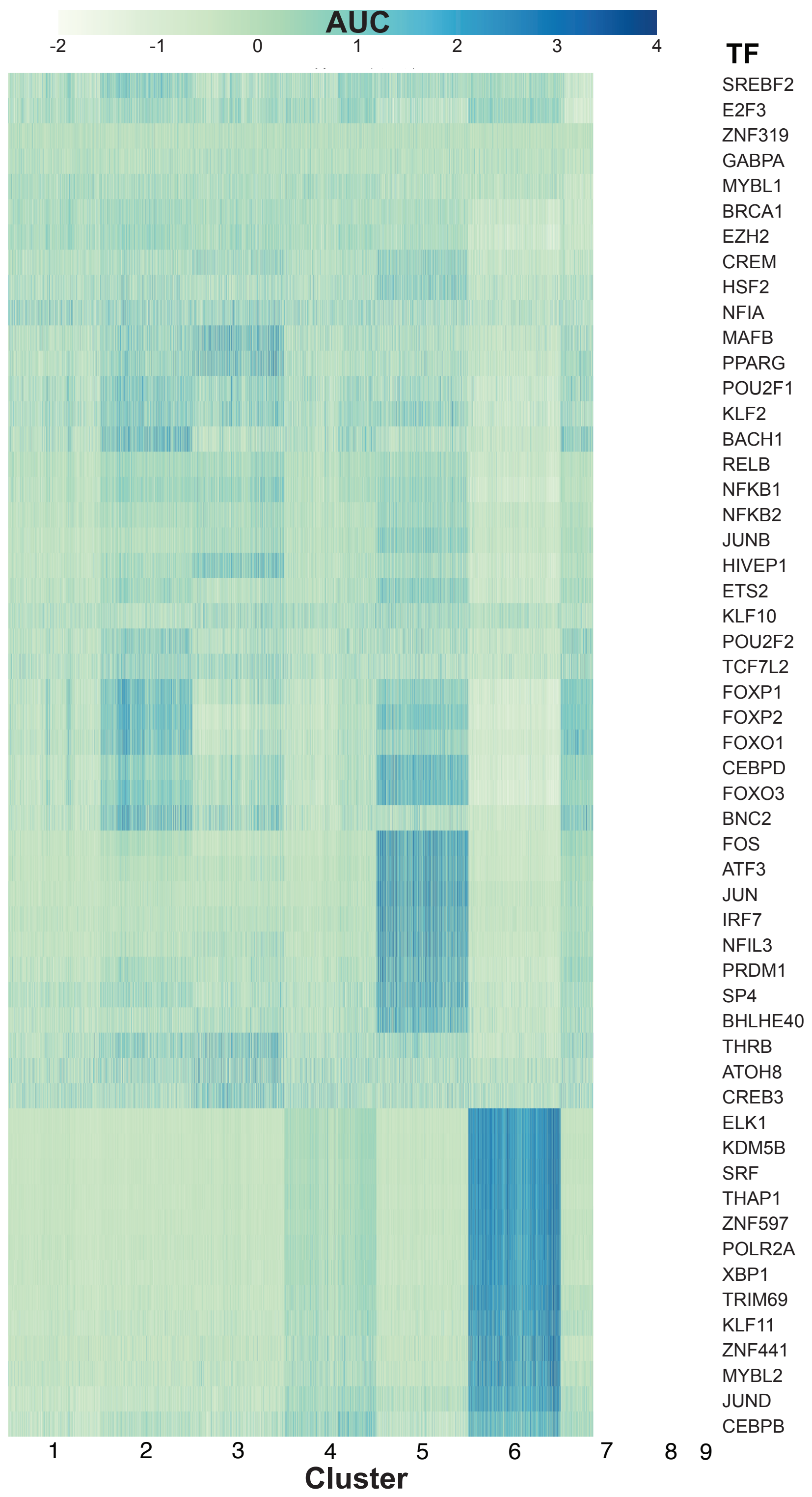


**Supplemental Figure 7. Transcription factor regulatory networks are specific and unique to subpopulations of microglia within the APOE ε3/ε3 genotype clusters.** Similar to the larger dataset, transcription factors driving gene expression within clusters are distinct when the data is comprised of only APOE ε3/ε3 allele carriers.

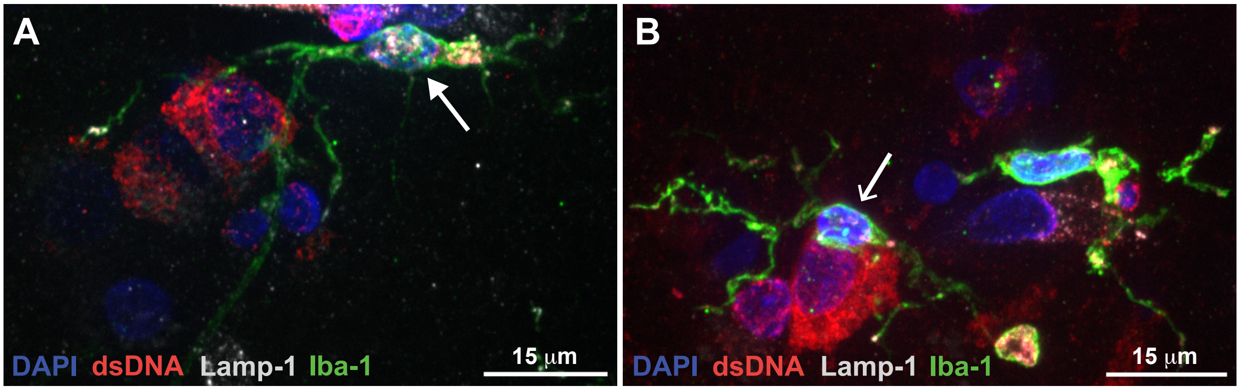


**Supplemental Figure 8. A subset of microglia demonstrate dsDNA signal and greater Lamp-1 signal in AD brain. A)** A representative example of an activated microglia with both large numbers of lysosomes (Lamp-1, white), and cytosolic dsDNA (red) in an AD case. **B)** Microglia (pointed arrowhead) without cytosolic dsDNA immunoreactivity (red) in the same case and tissue section appear ramified with less Lamp-1 signal.

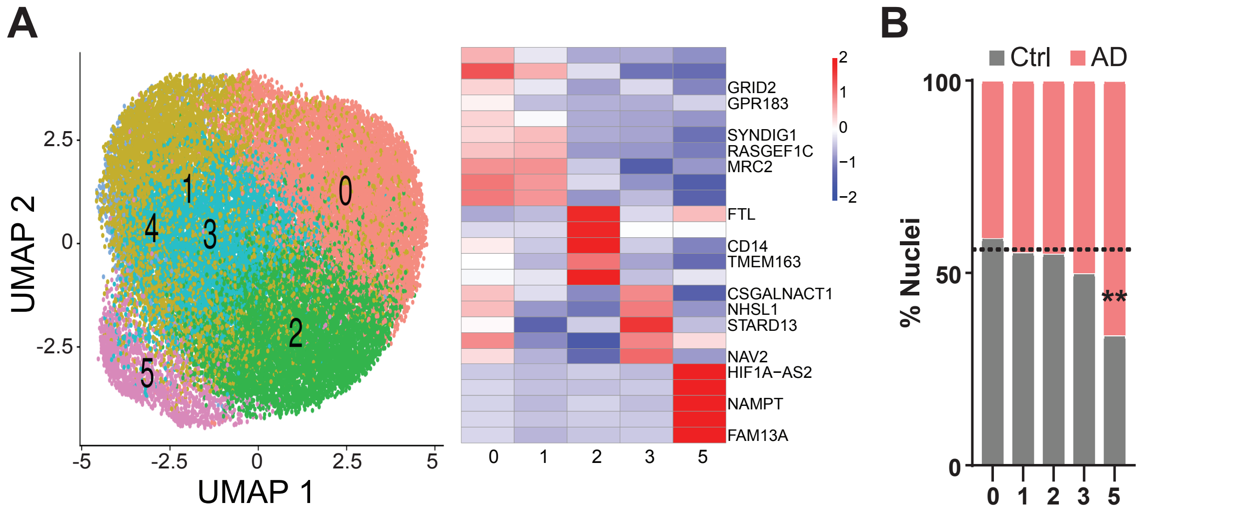


**Supplemental Figure 9. The APOE ε3/ε3 genotype cohort also demonstrates a subcluster of the homeostatic cluster increased in AD brain. A)** The results from the full dataset were replicated in the pure APOE ε3/ε3 allele cohort, suggesting multiple subpopulations exist within the "homeostatic" cluster. These subpopulations are distinct by gene expression despite being comprised of one "homeostatic" subpopulation. **B)** Within the APOE ε3/ε3 cohort "homeostatic" Cluster 1, there is a Cluster 1.5 that is significantly increased in AD brain like we found in the full cohort (Figure 6).

#### Table S4: Recipes for Nuclei Extraction and FANS Buffers:

**Myelin gradient buffer (Store at 4°C)**

| Reagent | Volume for 1 Liter |
|---|---|
| $NaH_2PO_4 \cdot H_2O$ (Fisher Scientific, S369-500), adjust to pH 7.4 with 3.56g/L $Na_2HPO_4 \cdot H_2O$ (Fluka, 71643) | 0.78 g |
| NaCl (Fisher Scientific, S271-3) | 8.0 g |
| KCl (Fisher Scientific, P217-500) | 0.4 g |
| Glucose (Sigma, G7021-1KG) | 2.0 g |
| BSA (VWR, EM-2930) | 0.2% |

**Nuclei buffer (NB) (Store at 4°C)**

| Reagent | Volume for 10mL |
|---|---|
| Nuclease-free water into a 15 ml conical tube. (Fisher Scientific, M46000) | 9.85 mL |
| 1 M Tris-HCl, pH 7.5 (ThermoFisher, 15567027) | 100 µL |
| 5 M NaCl (ThermoFisher AM9760G) | 20 µL |
| 1 M MgCl2 (ThermoFisher Scientific, AM9530G) | 30 µL |

**Nuclei lysis buffer (NLB) (make same-day)**

| Reagent | Volume for 1 Sample |
|---|---|
| Nuclei Buffer | 727 µL |
| 10% NP-40 alternative (final concentration 0.1%) | 10 µL |
| Protease inhibitors in DPBS (Sigma-Aldrich, 4693124001) | 142.9 µL |
| 1mM ATA in NB (make fresh evening prior) | 112.5 µL |
| PMSF (Tocris Bioscience, 4486) | 10 µL |
| Phosphatase inhibitors (Sigma-Aldrich, P5726-1ML) | 5 µL |
| Protector RNase inhibitor (final concentration 1 U/µl). (Sigma, 3335402001) | 28.25 µL |
| **Total Volume** | 1,035.5 µL |

**Nuclei suspension solution (NSS) (make same-day) (10X Genomics calls this "Wash Buffer")**

| Reagent | Volume for 1 Sample |
|---|---|
| DPBS (Sigma-Aldrich, D8537-500ML) | 637 µL |
| 10% BSA (final concentration 1%) (Sigma-Aldrich, A1595-50mL) | 100 µL |
| 1mM ATA in DPBS (make fresh evening prior) | 100 µL |
| 7x Protease inhibitors in DPBS (Sigma-Aldrich, 4693124001) | 142.9 µL |
| 1M PMSF (Tocris Bioscience, 4486) | 10 µL |
| Protector RNase inhibitor (final concentration 1.0 U/µl) (Sigma, 3335402001) | 5 µL |
| Phosphatase inhibitors (Sigma-Aldrich, P5726-1ML) | 5 µL |
| **Total Volume** | 1 mL |

**Percoll/myelin gradient buffer solution (PMB) (make same day)**

| **Reagent** | **Volume for 1 Sample** |
|---|---|
| Myelin gradient buffer | 412 µL |
| 10x HBSS (Fisher Sci., 14185052) | 30 µL |
| 1mM ATA in myelin gradient buffer (make fresh evening prior) | 100 uL |
| 1.5M NaCl | 25 µL |
| Percoll (Fisher Sci., 17-089-101) (Do not use if crystals have precipitated out) | 270 µL |
| Protector RNase Inhibitor (final concentration 1.0 U/µl) (Sigma Aldrich, 3335402001) | 5 µL |
| Phosphatase inhibitors (Sigma-Aldrich, P5726-1ML) | 5 µL |
| 7x Protease inhibitors in myelin gradient buffer (Sigma-Aldrich, 4693124001) | 142.9 µL |
| PMSF (Tocris Bioscience, 4486) | 10 µL |
| **Total Volume** | 1 mL |

**FACS Media (FM) (store at 4°C)**

| **Reagent** | **Volume for 1 sample** |
|---|---|
| Nuclease free $H_2O$ (Fisher Scientific, M46000) | 7.7 mL |
| HEPES (Invitrogen, 15630080) | 100 uL |
| 10x HBSS without Mg/Ca (Fisher Scientific, 14185052) | 1 mL |
| FBS 10% | 1 mL |
| Protease inhibitors (Sigma-Aldrich, 4693124001) (dissolve tablet in 9.8mL of the above media at room temp and chill on ice before adding the solutions below) | "~200 uL" |
| PMSF (Tocris Bioscience, 4486) | 100 uL |
| Phosphatase inhibitors (Sigma-Aldrich, P5726-1ML) | 50 uL |
| Protector RNase Inhibitor (Sigma Aldrich, 3335402001) | 50 uL |
| 100 uM ATA in dPBS (Sigma-Aldrich, A1895) good at -20C for 1 month | 1 mL |
| Total Volume | 11.2 mL |
